# Switching Promotor Recognition of Phage RNA Polymerase in Silico Following Path along Lab Directed Evolution

**DOI:** 10.1101/2021.07.22.453425

**Authors:** E Chao, Liqiang Dai, Jin Yu

## Abstract

In this work we computationally investigated how a viral RNA polymerase (RNAP) from bacteriophage T7 evolves into RNAP variants under lab-directed evolution to switch recognition from T7 promoter to T3 promoter in transcription initiation. We first constructed a closed initiation complex for the wild-type T7 RNAP, and then for six mutant RNAPs discovered from phage assisted continuous evolution experiments. All-atom molecular dynamics (MD) simulations up to one microsecond each were conducted on these RNAPs in complex with T7/T3 promoter. Our simulations show notably that protein-DNA electrostatic interactions or stabilities at the RNAP-DNA promoter interface well dictate the promoter recognition preference of the RNAP and variants. Key residues and structural elements that contribute significantly to switching the promoter recognition were identified. Followed by a first point mutation N748D on the specificity loop to slightly disengage the RNAP from the promoter to hinder the original recognition, we found an auxiliary helix (206-225) that takes over switching the promoter recognition upon further mutations (E222K and E207K), by forming additional charge interactions with the promoter DNA and reorientating differently on the T7 and T3 promoter. Further mutations on the AT-rich loop and the specificity loop can fully switch the RNAP-promoter recognition to the T3 promoter. Overall, our studies reveal energetics and structural dynamics details along an exemplary directed evolutionary path of the phage RNAP variants for a rewired promoter recognition function. The findings demonstrate underlying physical mechanisms and are expected to assist knowledge/data learning or rational redesign of the protein enzyme structure-function.

## Introduction

Lab directed evolution technologies in recent years have made substantial advancements in functional design or redesign of biomolecular systems, in particular, on protein enzyme activities and specificities [1-5]. In the lab directed evolution, sequence mutations and recombination are intensively promoted and followed by high-throughput screening or selection to target on specific protein functions [6-9]. For example, in phage-assisted continuous evolution (PACE), bacteriophage with modified life cycle is employed to transfer evolving genes between bacterial host-cells to promote fast replicating phage populations which contain gene mutations toward certain favored enzyme activities [10, 11]. With technology advancements, not only individual protein enzymes with certain functions can be designed, but also pathway or protein interaction network can be modulated or rewired [12-17]. Meanwhile, rational design of protein functions based on molecular structures and biochemical properties of the protein have always been pursued [18-22], which usually demand physical understanding and computational exploration on optimal solutions in the high-dimensional space of protein sequence or conformation evolution. Since biomolecules or enzymes are intrinsically complex systems evolved with highly complicated structure-function relations, straightforward physical or rational approaches are usually highly challenging. The lab directed evolution studies, however, provide abundant data on designed/redesigned on-path and end products with desired functions, which can be particularly interesting to learn and to infer the underlying structure-function relation, so that to further support physically based rational approaches. In this work, we use *in silico* methods, i.e., molecular modeling and all-atom molecular dynamics (MD) simulation to study promoter recognition of viral RNA polymerase (RNAP) variants discovered from lab directed evolution. In particular, we take advantage of the PACE achievements on switching a bacteriophage RNAP from recognizing its original promoter to another one in a similar phage system [9, 23].

The recognition and binding of RNAP to the promoter occur at the initial stage of gene transcription, which essentially determines the promoter activity or productivity of followed gene expression. In eukaryote cells, the transcription initiation is highly regulated, conducted by a multi-subunit RNAP in coordination with a large number of transcription factors [24, 25]. In contrast, single-subunit viral RNAP from bacteriophage T7, which is constantly utilized in lab gene expression system, is able to complete transcription from initiation to termination in the absence of additional factors [26, 27]. Such viral RNAP system is accordingly ideal for studying elementary key transcription functions. In particular, T7 RNAP and its closely related single-subunit viral RNAPs from other bacteriophages (e.g. T3, SP6, K11) demonstrate high specificities in their individual promoter activities [28-31]. Meanwhile, mutant phage RNAPs have also been identified to show modulated promoter specificities, e.g. certain mutations of T7 RNAP lead to switching of its specificity from T7 promotor to T3 or SP6 promoter [32-34]. Hence, study of specific promotor recognition in phage RNAP transcription initiation, in particular, on how the wild-type (or wt) RNAP and its mutant (or mt) RNAPs or variants change their promoter specificities from the original DNA promotor to the promoter of alternated DNA sequences, would be of high interest to reveal physical mechanisms underlying specific protein-DNA sequence recognition.

In order to study the phage RNAP promoter recognition using in silico approach, high-resolution atomic structures of the corresponding systems are needed. The high-resolution crystal structure of the T7 RNAP-DNA promoter binding complex has been resolved [35], with the RNAP in association with an incomplete transcription bubble that demonstrates an open form (see **Fig 1A**). However, in the stage of RNAP initial binding and recognition on the promotor, the promoter DNA still remains closed, hence a closed promotor complex of T7 RNAP is needed for this study. Employing molecular docking and modeling techniques, we constructed a closed transcription initiation complex of T7 RNAP (see **Fig 1B**), so that the specific binding characteristics of T7 RNAP to its promoter can be directly investigated by using all-atom molecular dynamics (MD) simulations. Subsequently, *in silico* mutations of a small number of protein residues (up to 5-6 mutations) were conducted to the wt-RNAP to obtain the RNAP variants, which have then been studied further via the all-atom MD simulations.

**Fig 1.**
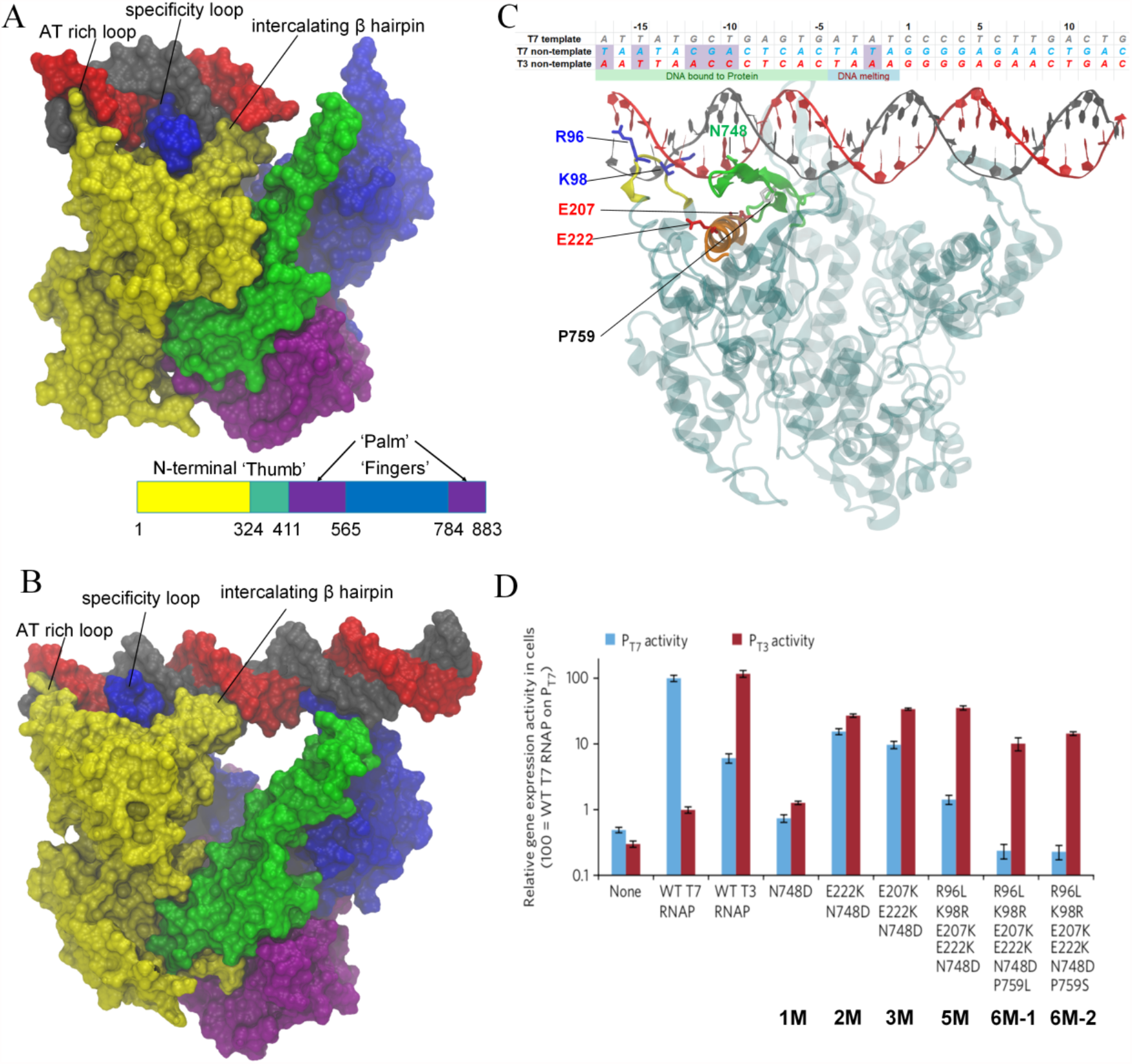
Structures of the closed and open complexes of T7 RNAP during transcription initiation and mutant RNAPs from recognizing T7 DNA promotor to T3 promotor identified from directed evolution experiment [23]. (A) The crystal structure of the T7 RNAP taken from the open complex (PDB code: 1QLN) [41], in the absence of transcript bubble. (B) The constructed model of the closed T7 RNAP initiation complex in this work. (C) The T7 RNAP promotor recognition or the closed initiation complex, with amino acids mutated in the directed evolution shown. The AT-rich loop (ATL, residue 93-101, yellow), specificity loop (SPL, residue 739-770, green), intercalating β hairpin (INB, residue 230-245, pink), and an auxiliary helix (AXH, residue 206-225, orange) are shown. The protein is shown in transparent cyan. The template and non-template T7 DNA promoter strands are shown in gray and red, respectively, with corresponding sequence listed for both T7 and T3 promoter (differences shaded). (D) The gene expression activities of T7 RNAP variants containing subsets of mutations (labeled 1M to 6M) in the evolved clones from the PACE, working on the T7 (blue bars) and T3 promoters (red). The data show mean values ± s.e.m. The images are adapted from the experimental work [23].

Earlier studies show that the T7 DNA promoter is mainly composed of two functional domains, a protein binding region upstream (from -17 to -5) and a transcription initiation region downstream (from -4 to +6) (see **Fig 1C**) [36]. In general, experiments found that the substitution of bases in the upstream binding region has significant impact on the RNAP binding [37, 38], but little impact on the initiation; the replacement of bases in the downstream region, however, mainly affect the initiation, but not the RNAP binding [31, 39]. In addition, studies have shown that the DNA region responsible for specific binding and recognition in phage T7 promoter is largely via -12 to -8, while the region distinguishes the T7 and T3 promoter locates mainly around -12 to -10 [38, 40].

In the PACE experiments system [9, 23], the wt-T7 RNAP that recognizes T7 promoter is evolved into variant RNAPs that can finally recognize the promoter from phage T3, with their corresponding promoter activities documented [23] (see **Fig 1D**). At an initial evolution stage, the wt-T7 RNAP can transcribe from the T7 promoter but not from the T3 promoter. Next, a single-point mutant (or 1M: N748D) appears, which recognizes neither the T7 nor the T3 promoter, as both promoter activities are low. Further, a double mutant (or 2M: E222K & N748D) and a triple mutant (or 3M: E207K, E222K & N748D) show increased promoter activities on both T7 and T3 promoter, with activities on the T3 promoter slightly higher than that on the T7 promoter. Finally, a five-point mutant (or 5M: R96L, K98R, E207K, E222K, & N748D) and two six-point mutants (6M-1 or 6M-2: R96L, K98R, E207K, E222K, N748D, &P759L or P759S) are evolved, which are significantly active on the T3 promoter but function marginally on the T7 promoter, demonstrating prominent bias toward recognizing the T3 promoter. Note that all involved amino acid mutations are located at an AT-rich loop (ATL; residue 93-101), an auxiliary helix (AXH; residue 206-225), an intercalating beta hairpin (INB residue 230-245), and a specificity loop (SPL; residue 739-770), which are all located at the RNAP-promoter DNA binding interface or nearby (see **Fig 1C**) [41]. The ATL and SPL had been previously recognized as the key structural elements for promoter recognition in T7 RNAP [41, 42], while the mutations on the AXH were discovered from the directed evolution experiment system [9, 23] and are particularly analyzed in this work. All these mutations are accordingly studied in current *in silico* investigations, with seven RNAPs (wt T7 RNAP and six variants) modeled and simulated at the atomic resolution, and each RNAP in complex with T7 and T3 promoter, respectively.

By following the phage T7 RNAP protein and its variants on the lab directed evolution path obtained from the PACE, rewired toward recognizing the phage T3 promoter, we intend to understand how the protein-DNA recognition is achieved, specifically, and how the specific recognition function is modulated via point mutations, under pressure of the directed evolution. To do that, we conducted all-atom MD simulations up to one microsecond for each of the above RNAP-promoter DNA complexes. We comparatively studied protein-DNA interactions at the promotor recognition sites for all these systems. We found that electrostatic interactions between the RNAP protein and DNA provide an effective measure to highlight the protein-DNA recognition preference in current systems. Certain residues seem to play key roles in switching the promoter recognition function of the protein. In addition to the ATL and SPL structural elements previously identified, the AXH element turns out to contribute significantly to switch the promoter binding and recognition. In particular, the coordination and competition among these essential structural elements to the promotor DNA are also examined via hydrogen bonding (HB) and salt-bridge (SB) interactions.

## Results

### Protein-DNA electrostatic interaction energetics provides quantitative measures for the RNAP-promoter recognition

In order to probe whether protein-DNA interactions that stabilize the RNAP at the promoter also contribute to the promoter recognition and differentiation, we calculated the electrostatic (ele, *E*^*ele*^) and van der Waals (vdW, *E*^*vdw*^) interaction energies between the RNAP protein and the promoter binding region of the DNA (−17 To -5), for wt T7 RNAP and all mutants (14 simulation systems). The interactions were calculated between atoms from protein and DNA at a cutoff distance ∼25 Å (the results converge for cutoff > 20 Å, see **Methods** and **Supplementary Information** or **SI Fig S1A**). The convergences of the energetic calculations over simulation time are shown for the wt-RNAP (**Fig 2A**) and for the RNAP variants (see also **SI Fig S1B-F**). The energetics obtained by averaging one microsecond of simulation trajectories for individual systems are presented (**Fig 2B**).

**Fig 2.**
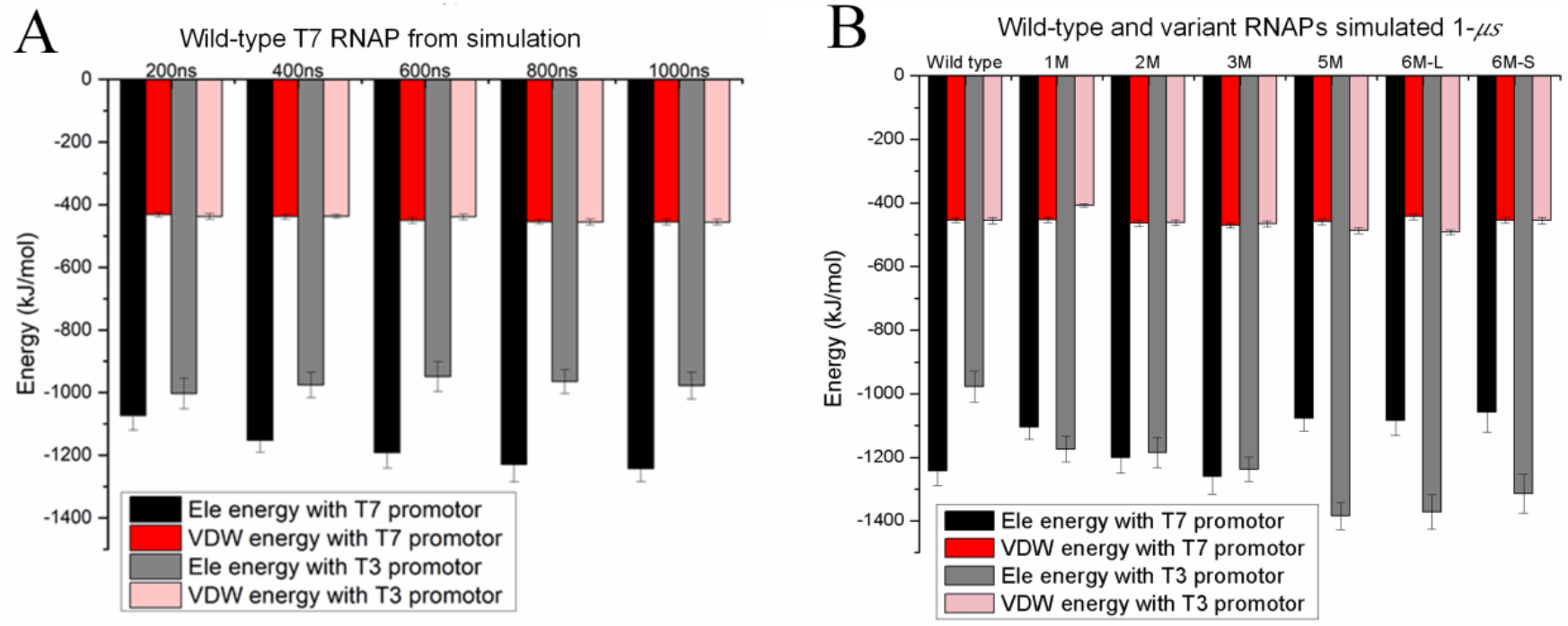
Protein-DNA promoter interaction energetics calculated from all-atom MD simulations for the wt T7 RNAP and six RNAP variants in the lab directed evolution. The interaction energetics include electrostatic (ele, *E*^*ele*^) and van der Waals (vdW, *E*^*vdw*^) contributions, averaged from individual simulations. (A) The interaction energetics between the wt-RNAP and T7/T3 promoter from simulations of 100 ns to 1000 ns or 1 μs in length. Convergence shows after ∼ 400 ns. (B) The interaction energetics averaged over the 1-μs trajectories for 14 simulation systems, i.e. wt-RNAP and six mt-RNAPs (1M to 6M; see **Fig S1** for simulation convergence), in complex with T7/T3 promotor.

The results indicate that the bias of *E*^*ele*^ between the RNAP and the promoter binding region well characterizes the recognition preferences of the RNAP, as being shown in the experimental results (**Fig 1D**). For example, the wt T7 RNAP has lower electrostatic interaction energies *E*^*ele*^ with the T7 promoter than with the T3 promoter, i.e., it binds more stably and electrostatically to the T7 promoter, consistent with it having higher activities or recognizing better the T7 than T3 promoter. For 1M, 2M and 3M RNAP variants, they bind T7 and T3 promoters with similar *E*^*ele*^, consistent with their low differentiation in the promoter activities between the T7 and T3. Notably, for 5M and 6M-1/2 RNAPs that demonstrate high activities on the T3 but not T7 promoter, the protein-DNA electrostatic energetics is significantly lower *E*^*ele*^ on the T3 promoter than on the T7 promoter. Meanwhile, the vdW energetics also show a similar tendency in stabilizing the protein with the promoter of higher activities, though not as significant as the electrostatic energetics (**Fig 2**).

### Individual residue contributions to the RNAP-promoter electrostatic bias and corresponding dynamics at the protein-DNA binding interface

Since for certain RNAP, the protein-DNA energetic difference between the T7 and T3 promoter well characterizes the promoter recognition preference, we calculated 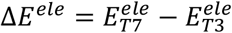 for each system (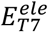 and 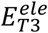 are the RNAP interaction energetics with the T7 and T3 promoters, respectively) and projected the contributions to Δ*E*^*ele*^ onto individual amino acids (AAs), as 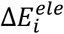 for the *i-th* AA from the RNAP (**Fig 3**; or see individual AA contributions to respective 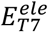 and 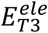 in **SI Fig 2**; numerical values in **Table S1**). In particular, we found that the key AAs contributing significantly to the promoter recognition, i.e., with large amplitudes of 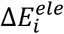, locate mainly on the ATL (AAs 93-101), AXH (206-225), INB (230-245), and SPL (739-770). For the wt RNAP, ATL-R96, AXH-R215, INB-R231, and SPL-R746/756 stabilize both T7 and T3 promoters (see **SI Fig S2**), more to T7 and less to T3; Q135 (located between INB and AXH) only interacts noticeably with T7. Correspondingly, ATL-R96, Q135, AXH-R215, INB-R231, SPL-R746 having 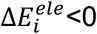, i.e., biasing the RNAP to be more stabilized on the T7 promotor (**Fig 3B**). In addition, SPL-N748 shows bias toward the T7 promoter, though its respective interactions with T7/T3 promoter are not particularly strong (**SI Fig S2**). AXH-E218 on the other hand, however, biases toward the T3 promoter (with 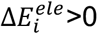), without noticeable interactions with the respective promoters either.

**Fig 3.**
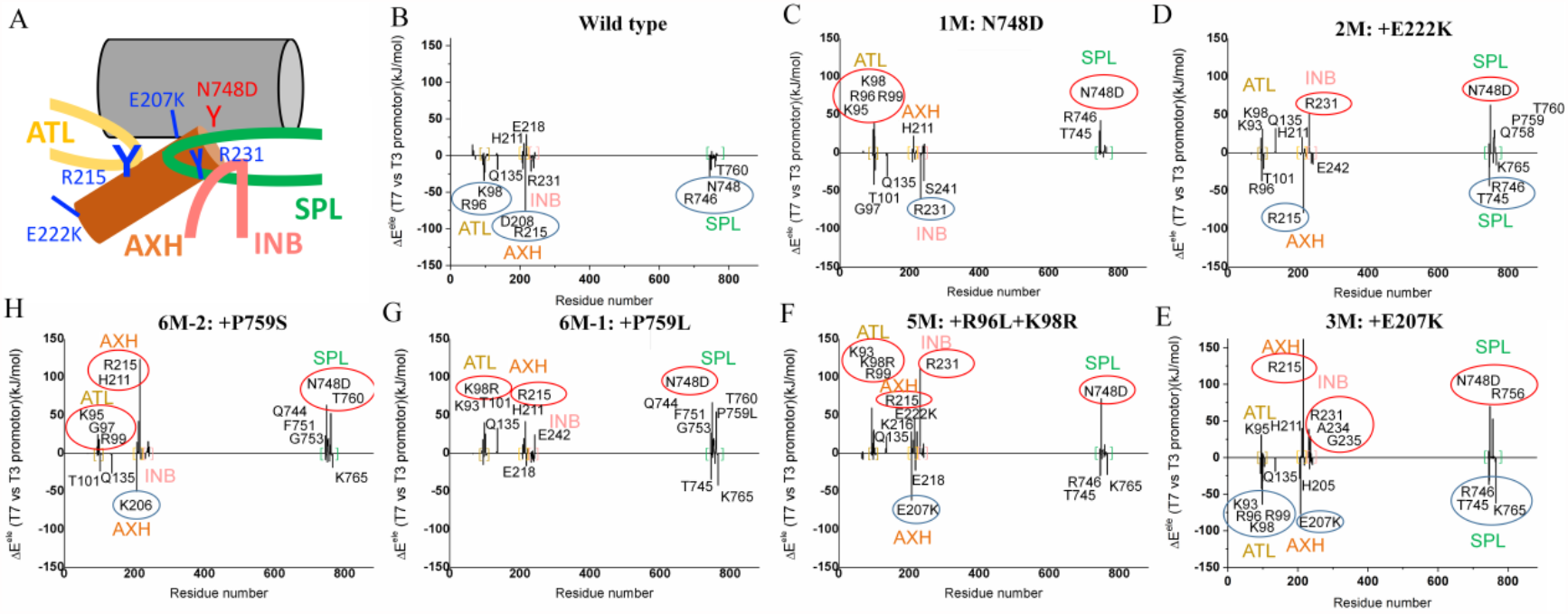
The energetic contributions 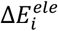 from individual amino acids that bias or stabilize the RNAP association with the T7 promotor 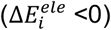 or with the T3 promotor 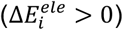, as 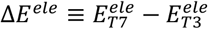 is the relative protein-DNA electrostatic interaction energy calculated between the RNAPs and the two promotors. The energetic contributions are demonstrated for the wt to mt-RNAPs (1M to 6M). Key residues with notable contributions to the energy bias are labeled (with |Δ*E*^*ele*^| > ∼ 15 *KJ*/*mol*).

In the 1M (N748D, **Fig 3C**), the mutation itself places an immediate energetic bias toward stabilizing the T3 promoter (it is indeed D748 in T3 RNAP) [43]. However, energetic contributions of N748D toward either T7 or T3 promoter is still insignificant (see **SI Fig S2**). Though ATL-G97 (&T101), Q135, and INB-R231 still having 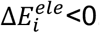, ATL-K95 & K98 (along with R96&KR99), AXH-H211 and SPL-N748D (along with T745&R746) start having notable 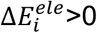, i.e., biasing toward the T3 promoter. The 1M variant thus has the SPL destabilization introduced directly via the N748D mutation, which then perturbs the promoter bias on both ATL and AXH (see **Fig 4A** and **B**).

**Fig 4.**
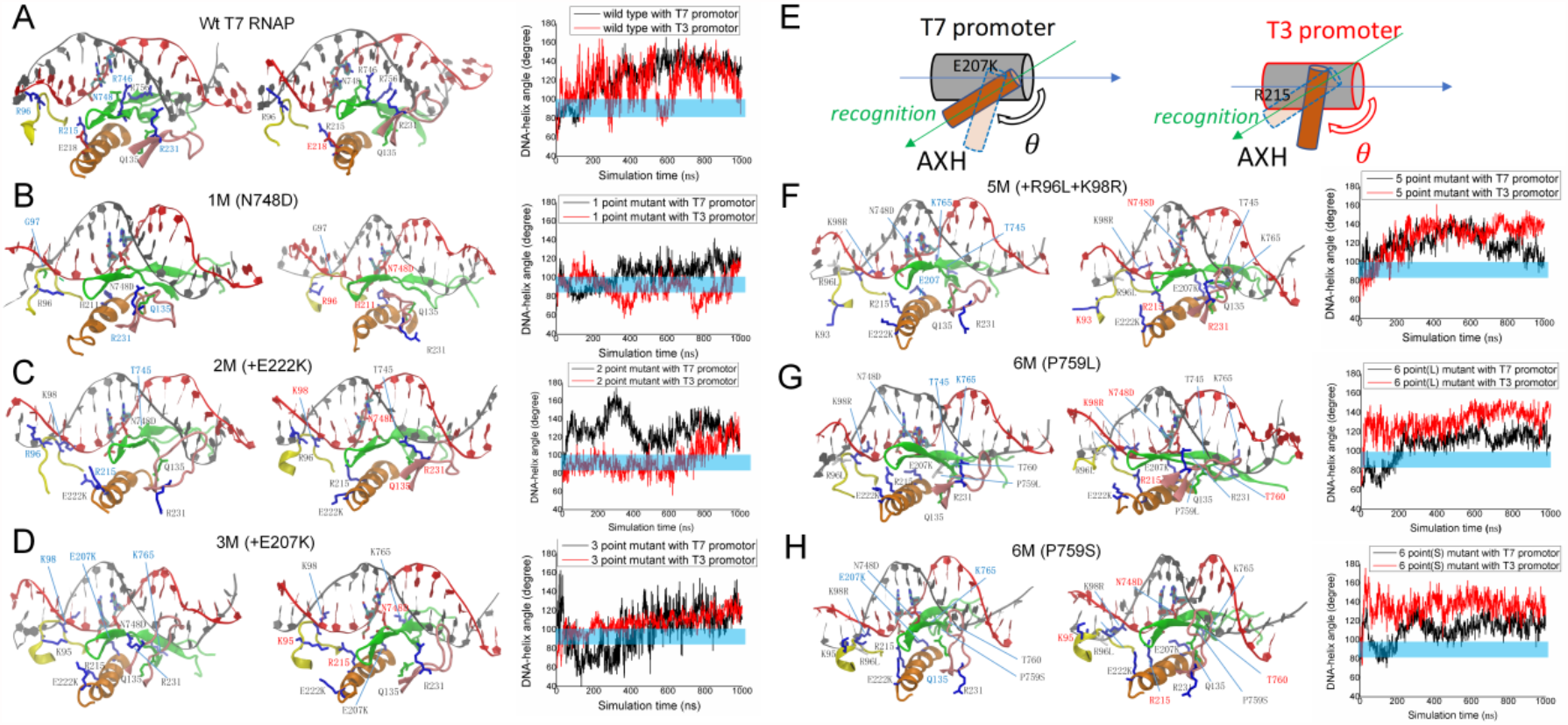
Structural views of the protein-DNA binding interface for wt-RNAP and six RNA variants (1M to 6M) in complexes with T7/T3 promoter. The AT-rich loop (ATL, in yellow), specificity loop (SPL, green), intercalating β hairpin (INB, pink) and auxiliary helix (AXH, in orange) in close association with the DNA promoter are shown. (A-D) Wt-RNAP and 1M-3M early mutants in the evolution. (F-H) RNAP mutants 5M to two 6M late in the evolution which recognize preferentially the T3 promoter. The orientation angle (θ) between the AXH and the DNA long axis are measured from the simulations (the time series to the right of the structural views for each system, with black/red data for T7/T3 promoter system, blue bars indicating vertical positioning of the AXH (θ∼90°) dis-advantageous to the RNAP promoter recognition. (E) A cartoon representation shows how the AXH orientation angle (θ) is measured, along with prominent angular changes of the AXH as well as E207K & R215 coordination (see text) on the T7 and T3 promoter DNA shown, respectively.

The 2M (N748D+E222K, **Fig 3D**) then has an additional AXH-E222K mutation. Now ATL-R96 (instead of G97) & T101, AXH-R215 and SPL-T745&R746 (along with K765) contribute 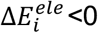, while ATL-K98, Q135, AXH-H211, INB-R231 and SPL-N748D (along with Q758&P759&T760) have 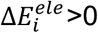. It seems that AXH-E222K mutation leads to the switch of the promoter preference for both Q135 and INB-R231 (see **Fig 4C**). Close examinations show that INB-R231 can switch its side chain up-side-down from 1M to 2M as E222K brings its side chain towards the T7 promoter (K222 still ∼10 Å away) but not the T3 promoter (see **SI Fig S3** for close views and analyses), accordingly, R231 associates much better with the T3 promoter than the T7 promoter in 2M.

Next, for 3M (N748D+E222K+E207K, **Fig 3E**) with a third mutation AXH-E207K, one obtains ATL-K98 (along with R96&R99), H205, AXH-E207K, SPL-T745&R746 &K765 having 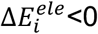 to stabilize the T7 promoter, and ATL-K95, AXH-H211&R215, INB-R231 (along with A234&G235) and SPL-N748D (along with R756) having 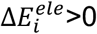, bias toward the T3 promoter. N748D has a significant stabilizing interaction with the T3 promoter (larger than that in 1M or 2M), and INB-R231 interacts closely with both T7 and T3 promoters (see **SI Fig S2**), yet maintains its bias toward T3. This time, although AXH-E207K itself largely stabilizes and biases toward the T7 promoter, it facilitates AXH-R215 to bind preferably toward the T3 promoter (but not the T7 promoter). A competition between E207K and R215 toward the DNA promoter actually shows, as E207K succeeds binding more closely than R215 to the T7 promoter, R215 actually binds more closely than E207K toward the T3 promoter (see **Fig 4D and SI Fig S4** for close views and analyses). Due to promoter interactions with E207K and R215 from N and C terminal of the AXH, respectively, the AXH orientation angle (θ) with the DNA long axis substantially changes (see **Fig 4D**), with θ_T7_ decreases from 130° ±15 ° (in 2M) to 104 ° ± 21 ° in 3M, and θ_T3_ increases from 97°±17° (in 2M) to 111° ± 8° in 3M. The θ_T7_ change thus brings the AXH more vertically aligned with the DNA, and the trend persists into 5M and 6M; an opposite trend shows for θ_T3_ (i.e., the AXH aligned better with the T3 promoter DNA axis then). Overall, the 2M and 3M do not differentiate much between the T7 and T3 promoter, yet they do prepare for the necessary or key reside configurations for the promoter recognition/differentiation in 5M/6M. In particular, R215 biasing toward the T3 promoter and the accompanied AXH orientational change with respect to the promoter DNA appear to be essential.

In comparison, the promoter recognition and differentiation become prominent in 5M and 6M, in which R96L+K98R on the ATL additionally occur (5M, **Fig 3F**), and then SPL-P759L/S (6M, **Fig 3G**&**H**). In both cases, there are more residues contributing to Δ*E*^*ele*^ > 0, i.e., to stabilize the T3 promoter. In 5M, AXH-E207K (along with E218) and SPL-T745 (along with R746&K765) remain for 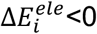, while ATL-K93 (along with K98R&R99), AXH-R215&E222K, INB-R231 and SPL-N748D all have 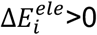. Since it is exactly K96 and R98 bias toward the T7 promoter in 2M and 3M, respectively, mutation of both largely abolishes the ATL bias on the T7 promoter (though K98R still closely interacts with both T7 and T3 promoter; see **SI Fig S2**). The SPL-K765 stabilization toward the T7 promoter also disappears comparing to 3M.

In the 6M-1 (P759L), SPL-T745 and SPL-K765 can still contribute to 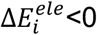, while ATL-K98R, AXH-R215&H211, SPL-N748D (along with several residues from Q744 to T760) have 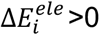. In the 6M-2 (P759S), ATL-T101, AXH-K206 and SPL-K765 contribute to 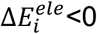, ATL-K95 to R99, AXH-R215&H211, SFL-N748D (and residues from Q744 to T760) have 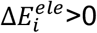. It seems that SPL-P759L/S mutation induces Q744 to T760 region to stabilize the T3 promoter further, while the flexible INB-R231 does not necessarily bias toward the T3 promoter (it interacts with the promoter closely with both T7 and T3 in 6M-1, see **SI Fig S2**). The AXH-E207K stabilization to the T7 promoter does not persist in 6M anymore. Marginal ATL/AXH/SPL stabilization to the T7 promoter still exists in the 5M or 6M. Meanwhile, SPL-N748D (started from 1M) and AXH-R215 (triggered in 3M) contribute robustly toward the T3 promoter.

### Analyzing hydrogen bonds (HBs) at the RNAP-DNA promoter interface to probe further AA contribution to recognition

In order to investigate the specific recognition of the RNAP on the promoter, we checked the corresponding hydrogen bonds (HBs) and salt bridge (SB) interactions at the RNAP-promoter interface for each of the simulation systems (see **Methods**). Since most of HBs are fluctuating and highly dynamical, we recorded HBs with at least ∼ 10% of the occupancy during the microsecond simulation (∼0.8 μs). The corresponding results are summarized in **Fig 5A**. Note that the DNA sequences of the T7 and T3 promoter differ only at -17, -15, -12, -11 and -10 positions (around the DNA major groove -15 to -10; see **Fig 1D**), and position -12 to -10 are crucial for the promoter specificity [36].

**Fig 5.**
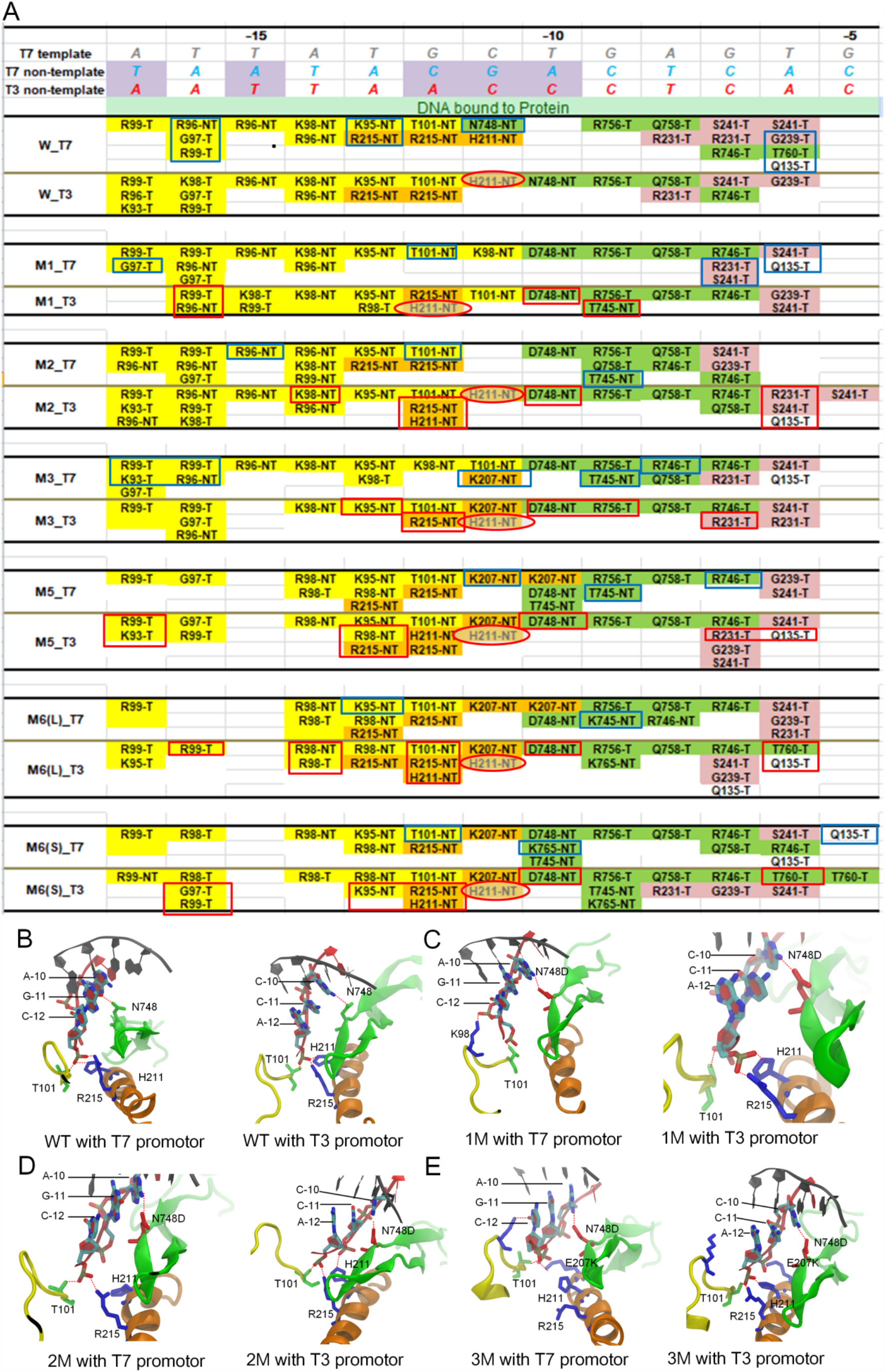
The hydrogen bond (HB) patterns between RNAP and the promotor DNA from the simulation systems. (A) The HB interactions formed (>10%) in the last 0.8 μs of the simulation between the RNAP and the bound region of the promotor DNA. The T7 DNA template (gray) and non-template (blue) chains are displayed schematically with sequences on top, with T3 non-template chain (red) shown as well (the different sequences between T7 and T3 promoters are highlighted in purple). The amino acids at the AT-rich loop (ATL), specificity loop (SPL), intercalating β hairpin (INB) and auxiliary helix (AXH) are highlighted by yellow, green, pink and orange, respectively. Blue and red rectangles are placed to show residues simultaneously contributing significantly in electrostatic stabilization (as in **Fig 3**) to the T7 and T3 promotors, respectively. The HB contributed from the AXH-H211-NT is circled red, as H211-NT always bias toward the T3 promoter in the mutant RNAPs. (B-E) The HBs formed between the RNAP and the central bound region (−12 to -10) of promotor DNA for the wt-RNAP (B), and for the early or transition mt-RNAPs 1M to 3M (C-E), respectively, on the T7 promoter (*left*) and T3 promoter (*right*).

The HBs formed by the ATL and DNA span extensively from the upstream minor groove region (−17 to -15) to the major groove in the middle (−15 to -12). In the wt-RNAP, ATL forms ∼4 HBs (with T7) and 7 HBs (with T3) at the upstream region (−17 to -15), mainly with the template strand (denoted T); one HB forms very downstream as T101: NT-12 (NT denotes the non-template strand), on both the T7 and T3 promoter. In the single-point mutant (1M), ATL forms ∼ 6 HBs (with T7) and ∼4 HBs (with T3) upstream; two HBs T101:NT-12 and K98:NT-11 are formed on the T7 promoter, and the two HBs switch to T101:NT-11 and K98:NT-12 on the T3 promoter. In the double and triple mutants (2M and 3M), ALT maintains ∼ 6HBs (with T7) but 4-7 HBs (with T3) upstream; the most downstream HB forms as T101:NT-12/-11 on the T7 promoter, or as T101:NT-12 on the T3 promoter. In the T3 promoter preferred mutants (5M and 6M), the upstream ATL HBs reduced significantly: 1-2 HBs for T7, and 3-4 HBs for T3; the most downstream HB is always maintained as T101:NT-12 no matter on which promoter. Hence, it seems that the ATL HB association with the upstream template DNA strand is less with T7 in the wt-RNAP, while the trend switches somehow in the transitional mutants (1M to 3M), and recovers a bit but with the overall ATL HB association with the promoter weakened in the directed mutants (5M and 6M, due to R96L and K98R mutations on the ATL). In particular, the ATL association most downstream T101:NT-12 can move to NT-11 in the transitional mutants (1M and 3M), but recovers to stable T101:NT-12 in the directed mutants.

As for the AAs on the AXH, all HBs are concentrated in the middle of the major groove on the non-template DNA strand (NT-13 to -11). In the wt-RNAP, AXH forms 3 same HBs (R215:NT-13, R215:NT-12 and H211:NT-11; see **Fig 5B**) with both T7 and T3 promoter. Hence, the HBs do not seem to contribute to promoter differentiation. In 1M, AXH loses the HB association with the T7 promoter entirely, while two AXH HBs (R215:NT-12 and H211:NT-12) maintained with the T3 promoter (**Fig 5C**). R215 and H211 HBs recover somehow in 2M (upon E222K), with H211 persistently forming HB biasedly on the T3 promoter, as well as in 3M (upon E207K and K207:NT-11 formed for both promoters; **Fig 5D** and **E**). K207 continues to associate with NT-11 or even NT-10 into the directed mutants (5M and 6M), on both T7 and T3 promoters, with R215 forming HBs with NT-13/-12 on both promoters, H211 remains associating preferentially with the T3 promoter (5M and 6M; see **SI Fig S5**). Hence, it seems that 1M (or SPL-N748D) critically breaks a balance of the AXH HB association with the promoter DNA between the two species (T7 and T3), enables AXH-H211 to maintain HB with the T3 promoter but not with the T7 promoter anymore; then the mutations E222K and E207K, i.e., directly emerge on the AXH, enhance the AXH association with the DNA, as well as support the HB preference of H211 to the T3 promoter, persistently into the directed mutants.

The HBs formed by the SPL and the promoter DNA are mainly located far downstream (mainly on template T-10 to -7), up to NT-11 (only for N748 in the wt-RNA with T7), or down to -6/-5 (T760 in the wt-RNAP with T7 or in the 6M with T3). Notably, N748D emerges as a first and most critical mutation, moves the base specific HB N748:NT-11 with T7 and N748:NT-10 with T3 in the wt-RNAP to D748:NT-10 for ALL the RNAP variants (from 1M to 6M; see **Fig 5** and **SI Fig S5**) . D748 is the only residue forms a HB with the DNA base (NT-10) in the transitional mutants (1M-3M) and into the directed mutants (5M-6M), on both T7 and T3 promoter; NT-10 associates with E207K and/or T745 additionally, only on T7 but not T3 promoter. Hence, N748D seems to be essential to re-position HB interaction for the promoter recognition and support the AXH for the re-wired promoter differentiation. Other HB forming residues in the SPL seem to maintain stable contacts in all systems, including the wt-RNAP and variants (R756:G-9, Q758:A-8, R746:G-7). Note that T760:T-6 exists in the wt-RNAP with the T7 promoter preference; it then switches to the T3 promoter preference (6M), due to the mutation P759L/S.

Finally, the INB region forms HBs mainly via R231, Q239, and S241 with template -7 and -6 position (occasionally with R231 to -8 or S241 to -5). In the wt-RNAP, INB forms a couple of more HBs with the T7 promoter than with the T3 promoter. Such a bias reduces or even reverses slightly in the RNAP variants. It appears that INB can play some role still. In particular, INB-R231 shows a transient role in promoting bias toward the T3 promoter (in 2M and 5M), energetically or via forming HBs with the DNA, yet in general R231 side chain is highly flexible and frequently swings, without sustained bias.

In addition, we also examined the SB interactions at the RNAP-promoter DNA interface (see **SI Fig S6**). ATL at upstream contribute dominantly to the SB interactions, which shows no obvious differentiation between T7 and T3 promoter. Nevertheless, in the wt-RNAP, the ATL SB R96-NT-16 with the T7 promoter does contribute to energetically stabilize the T7 system; in the 2M/3M, the SB K98-T-15/T-14 on the T7 promoter does as well (see **Fig 2B**). Interestingly, such stabilization and bias abolish in the 5M/6M with R96L and K98R, which indicate that mutations on the ATL exactly promote the specificity to the T3 promoter. Meanwhile, at the -12 to -10 region key for the promoter differentiation, there are SBs from the AXH (e.g. R215-NT for all RNAPs except for M3 with T7; K207-NT starting from 3M), from ATL (K98 or R98 in all mt-RNAPs except for M3 with T7; occasionally K95-NT, in M1 with T3 and M3 with T7), and from SPL (occasionally K765-NT, for 1M/5M/6M with T7). In particular, one can see that the AXH involved with more SB interactions with DNA starting from 1M-2M upon the mutations, and the AXH-SBs extend further in 3M-6M.

## Discussion

The promoter binding of an RNAP plays a primary regulatory role in gene transcription. Since T7 RNAP can conduct initiation without transcription factor, it is expected that the RNAP can also search and locate the promoter, possibly via 1D diffusion along DNA, as detected experimentally [44]. Indeed, we tested the apo T7 RNAP search on the DNA non-specifically, using coarse-grained modeling and MD simulation [45]. We found that T7 RNAP diffuses processively along DNA with the SPL and the ATL structural elements making particularly frequent contacts with DNA (see **SI Fig S7**), due to protein-DNA electrostatic interactions (with an implicit solvent modeled at an ionic strength 0.15 M). Hence, it seems that the SPL and ATL can be the most important structural elements for the RNAP to locate the promoter sequences for initiation as the RNAP conducts diffusional search nonspecifically along DNA.

The directed evolution experiment was designed to train the RNAP from recognizing the T7 promoter to recognize the T3 promoter instead, and demonstrate a representative evolutionary path following the wt-RNAP →1M (N748D)→2M (+E222K) →3M (+E207K)→5M (+R96L&K98R)→6M-1 (+P759L) or 6M-2 (+P759S). Based on the promoter activities and differentiation, one can divide these RNAPs into four groups: the wt-RNAP, which recognizes the T7 but not T3 promoter; 1M, low promoter activities on both promoters; 2M and 3M, notable promoter activities on both promoters yet no differentiation; 5M and two 6Ms, which recognize the T3 but not T7 promoter. We analyzed the mechanism of switching the specific protein-DNA recognition along the above directed evolution path by conducting MD simulation of individual RNAPs along the path. According to previous experimental work [38] and the alignment of the T7 and T3 promoter sequence, it is noted that the -12 to 10 region of the promoter DNA mainly determines the specific sequence recognition. In particular, N748 from the SPL forms the only specific HB contact with the DNA base NT-11G (or NT-10C) on this region to the T7 (or T3) promoter. Accordingly, one expects that the HB between residue 748 and NT-11 or NT-10 is key to the specific recognition. Meanwhile, in the wt-RNAP, ATL extends from upstream to form HB contact T101:NT-12 downstream while AXH competitively binds -13 to -11 region (R215:NT-13&-12 and H211:NT-11) similarly on both T7 and T3 promoters. Followed, one sees that four critical protein-DNA binding/recognition transitions along the directed evolution path that play important roles.

The first mutation N748D from SPL breaks up the original binding and specificity from the wt-RNAP as transiting to the 1M RNAP. The mutation N748D indeed shifts the specific HB contact from NT-11G to NT-10A on the T7 promoter, while there is no shift on the T3 promoter (N748D:NT-11C maintains). In accompany, K98 and T101 from the upstream ATL extend from (NT-14 and NT-12 originally) to NT-11 downstream, on the T7 and T3 promoter, respectively. It appears as if ATL ‘pulls’ on the -12∼-10 promoter DNA toward upstream (see **Fig 6** schematics). AXH then behaves in response as abolishing R215&H211 HBs with NT-13 to -10 on the T7 promoter, while it adjusts the R215&H211 HBs from NT-13∼ -11 to NT-12 altogether on the T3 promoter. One thus sees that the ‘deactivation’ of the 1M RNAP on the T7 promoter simply results from perturbing some critical HBs (SPL-N748D, AXH-R215&H211). The consequent energetic impacts are noticeable, as the RNAP-T7 promoter interaction is destabilized while RNAP-T3 promoter interaction stabilized, electrostatically (the vdW still stabilizes or biases toward T7, see **Fig 2B**), with ATL-K95, AXH-H211 and SPL-N748D stabilizing toward T3.

**Fig 6.**
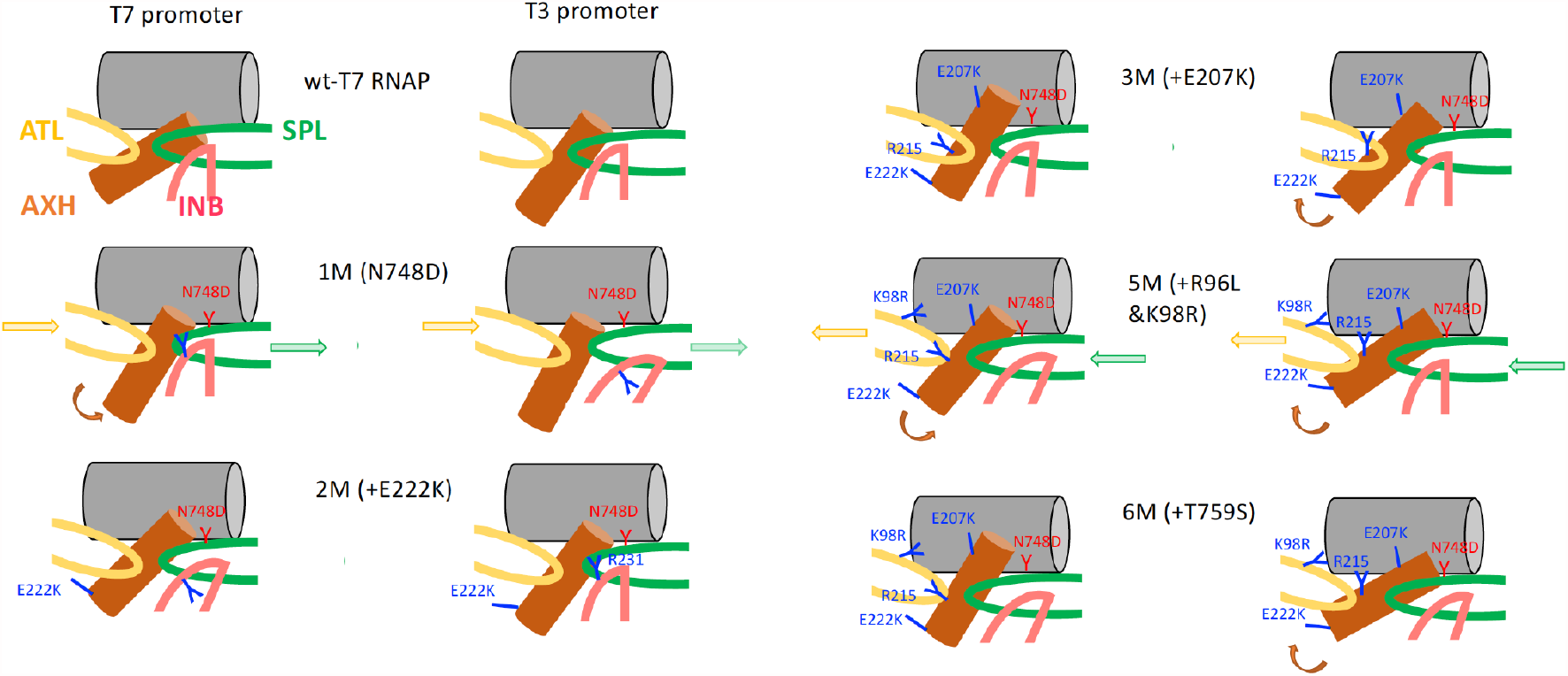
Schematics on the key structural elements/residues at RNAP-promoter DNA interface along the directed evolution path of switching the RNAP from recognizing the T7 promoter to recognize the T3 promoter. The key structural elements ATL (AT-rich loop, yellow), AXH (auxiliary helix, orange), INB (intercalating beta hairpin, red), and SPL (specificity loop, green) are presented along with most key residues in mutation and response. In the wt-RNAP to 1M transition (N748D bias toward T3), ATL and SPL shift downstream, AXH rotates (to be more vertical) on the T7 promoter. Upon 1M→ 2M (+E222K), AXH moves toward the DNA promoter slightly and starts to play a bigger role, while INB fluctuates somehow to allow transient R231 stabilization toward the T3 promoter. Upon 2M→ 3M (E207K), AXH-R215 moves toward the T3 promoter to align AXH better the DNA axis, while R215 cannot compete well with E207K on the T7 promoter. Upon 3M→ 5M (+R96L&K98R), ATL loses the biased R96&K98 interaction with T7; the ATL/SPL withdraws/extends upstream, so that AXH becomes even better aligned with the T3 promoter DNA but less with the T7 promoter DNA, which supports the switched promoter recognition to T3. 5M→ 6M (+T759S) further locks the T3 promoter specificity.

Next, AXH starts to play a bigger role via mutation E222K from 1M to 2M RNAP. The mutation of the negatively charged E222 on the AXH to the positively charged K222 allows the AXH to move closer to the promoter DNA (for both T7 and T3) than in the 1M. Consequently, AXH recovers its HBs with the T7 promoter via R215:NT-13&12, which also energetically stabilizes the T7 promoter binding. H211 additionally forms HB with NT-11 (aside with NT-12 in M1) on the T3 promoter. Energetically, E222K also triggers an immediate orientational switch of the side chain R231 from the INB, allosterically (e.g. the distance between two residues is about 15-22 Å), taking advantage of the flexible side chain motions of R231 on the INB-loop region so that to transiently stabilize R231 association with the T3 promoter. As R215 and R231 energetically bias toward the T7 and T3 promoter, respectively, there is no obvious energetic bias (either electrostatic or vdW) to T7/T3. 2M RNAP (N748D+E222K) accordingly shows notable promoter activities but does not display bias or differentiation between two promoters.

Then, AXH takes over the promoter binding via E207K from 2M to 3M. The additional charge conversion from negative to positive on the AXH allows it to be even closer to the promoter than 2M. E207K indeed well stabilizes energetical association of the 3M RNAP with the T7 promoter, and it also forms HB with NT-11 on both T7 and T3 promoter. Interestingly, however, the responses of R215 and consequent competition between K207 and R215 in association with the promoter reveal differently in the T7 and T3 systems. On the T7 promoter, R215 stays far from the DNA promoter and K207 dominates the energetic association and forms HB with NT-11; on the T3 promoter, however, R215 stays similarly close to the DNA promoter as K207, and they form HBs with NT-12 and NT-11, respectively. Thus, residue R215 originally favors the T7 promoter association (in the wt-RNAP) then switches to stabilize more or bias toward the T3 promoter upon the E207K mutation, and such R215 bias toward T3 then maintains robustly along the followed evolution path. One sees that upon the second and the third mutations, it is mainly the local electrostatic charge interactions that compete for the RNAP-promoter association, even though the overall energetic contributions between the two promoter systems are still in balance and show no bias.

Further, the ATL modulation via R96L+K98R fully switches the promoter bias to T3, transiting from 3M to 5M (and 6M). Although the most key residues N748K and R215 (enabled by E207K) energetically biasing toward the T3 promoter have established robust association, the further energetic stabilization to enable the promoter specificity is achieved by the ATL. Before, ATL remains energetically stabilizing to the T7 promoter and R96/K98 contributes to that, e.g. via the salt-bridge interactions. While R96L simply reduces the ATL-promoter association due to loss of electrostatic attraction, K98R abolishes HB with NT-12, so that ATL withdraws upstream on the T7 promoter, having T101 HB shifted from NT-11 (in 3M) to NT-12 (in 5M). Meanwhile, SPL-T745 extends upstream to form HB with NT-10 (along with D748) on the T7 promoter. Such changes may accordingly destabilize the AXH association with the T7 promoter. On the other hand, AXH seems to associate with the T3 promoter more extensively as H211 forms an additional HB with NT-12 (aside from NT-11), and R215 occasionally forms an additional HB with NT-13 (aside from NT-12), and aside from its robust energetic bias toward T3. With another mutation P759L/S (6M-1 or 6M-2) from the SPL, small energy bias from T760 to the T3 promoter is brought about, the energetic competition of K207 (weakening the bias toward T7) and R215 (strengthening the bias toward T3) on the AXH can be further tuned to bias toward the T3 promoter. Hence, the last stage enabling the promoter specificity seems to be achieved by further balancing a variety of HBs and local charge interactions.

## Conclusion

We utilized all-atom MD simulations to reveal physical mechanisms of viral T7 RNAP variants rewiring promoter recognition along the lab directed evolution path, as the promoter recognition of the RNAP switches from the original T7 promoter to the slightly different T3 promoter. As the first point mutation N748D emerges on the SPL (specificity loop) of T7 RNAP to bias toward the T3 promoter, it critically perturbs the HB patterns at the protein-DNA interface, shifts the balance between the SPL downstream and the ATL (AT-rich loop) upstream, and slightly dissociates the RNAP from the promoter to hinder the original promoter recognition. Notably, current study identifies an auxiliary helix (AXH 206-225) that takes over switching the RNAP-promoter recognition via the second and third mutations (E222K and E207K) of the RNAP along the directed evolution path, as AXH interacts more closely with the promoter mainly via the charge interactions upon the two mutations, and then reorientates differently on the T7 and T3 promoter to support further differentiation. The promoter specificity is finally switched upon mutations on the ATL (R96L+K98R), which adjust the protein-DNA HB and SB patterns and resets the balance between the ATL and SPL. Additional mutation on the SPL (R759L or R759S) modulates the RNAP-promoter interactions further and maintains the promoter specificity. Such structural dynamics and energetic details revealed from the simulations may assist structure-function information learning of the system to promote further rational design on specific RNAP-promoter recognition.

## Acknowledgements

This work has been supported by NSFC Grant #11775016 and #11635002. JY has been supported by the CMCF of UCI via NSF DMS 1763272 and the Simons Foundation grant #594598 and start-up fund from UCI. We also acknowledge the computational support from the Beijing Computational Science Research Center (CSRC).

## Materials and methods

### Obtaining an apo T7 RNAP protein structure

From a crystal structure of T7 RNAP (PDB:1ARO) [46] that contains additionally a T7 lysozyme (see **SI Fig S8**), we removed the lysozyme and used MODELLER[47] to fill in the missing gaps (residue id 60 to 72,165 to 182,234 to 240,345 to 384,590 to 611)in the protein. The obtained structure was then compared with an apo T7 RNAP structure containing Cα atoms only (PDB: 4RNP) [48] (see **SI Fig S8**), and consistency between the two is found.

### Docking the apo T7 RNAP onto double-stranded (ds) DNA promoter to construct a closed initiation complex

Using 2.0 version of web 3DNA (w3DNA 2.0) Interface[49], we generated standard B-form dsDNA containing the T7 promoter, with the template strand consisting of 30 nucleotides (3’ -ATTATGCTGAGTGATATCCCCTCTTGACTG-5’).

Then using Hdock Server [50], an online software for protein-protein and protein-DNA/RNA docking based on a hybrid algorithm of template-based modeling and *ab initio* free docking (http://hdock.phys.hust.edu.cn/), we docked the apo T7 RNAP structure onto the 30-bp dsDNA containing the T7 promotor. First, 100 complex structures generated from Hdock first. Next an FFT-based global docking program (HDOCK lite) was used to globally sample putative binding modes in the HDOCK server, in which an improved shape-based pairwise scoring function has been used, and the best scored top 10 structures were provided (as shown in **SI Fig S9**). From the top ten scored models, we selected the three structures which show similar DNA promoter positioning to that from the crystal structure of the T7 RNAP open initiation complex [41], then performed a 1-μs all-atom MD simulation, and calculated the HBs between the SPL and promoter in the structure. Finally, we selected the highest scored structure, which has protein-DNA HB interactions well represented, according to the existing open initiation complex structure of T7 RNAP (see **SI Fig S10**).

### Construction the coarse-grained (CG) model and setup of the CG simulations

The CG simulations were performed by the CafeMol 3.0 software [45]. The initial structure of T7 RNAP was obtained from the crystal structure (PDB ID: 1ARO) [47]. The CG protein structure was constructed by using the off-lattice Go model [51], each CG particle was located on the Cα atom to represent one amino acid, and with the conformations biased toward the native structure (crystal structure here) under the Go-model potential.

In the CG model of dsDNA (200 bp in length), each nucleotide is represented by three CG particles corresponding to base, sugar and phosphate groups via the 3SPN.1 model[51, 52], in which the bond stretching, angle bending, dihedral angle twisting, base-base interaction, excluded volume effect, solvation energy and electrostatic energy are considered. The electrostatic interactions and excluded volume effects are considered. All CG simulations are performed by Langevin dynamics under constant temperature with Berendsen thermostat.

### Construction of structural models of mutant RNAPs from directed evolution

According to the lab directed evolution (see **Fig 2**) [23], the wt-T7 RNAP gradually evolved to a series of mt-RNAPs that recognizes less the T7 promoter but more the T3 promoter. Based on those mutants, we constructed 14 simulation systems with 7 types of T7 RNAPs, including the wt-RNAP and variants (1M to 6M), and 2 types dsDNA (containing T7 or T3 promoter). For the mutation, The Tleap method in AmberTools is used to change the amino acid in the protein [53]. AmberTools is also used to mutate the DNA base pairs from the T7 promoter to those in the T3 promoter, and keeping the nucleic acid backbone unchanged. All these constructed structures are subjected to substantial energy minimization (20,000 steps energy minimization was conducted), and then perform the following MD simulations.

### Setup of atomistic MD simulations

All MD simulations were performed using GROMACS-5.1 software package [54-56]. The AMBER99sb-2012 force field with PARMBSC0 nucleic acid parameters [57-59] was used to describe the system. The minimum distance from the protein to the border of the simulation box was set to 13 Å. In order to neutralize the system and keep the ion concentration at an ionic strength of 0.15M, 163 Na^+^ ions and 119 Cl^-^ ions were added. The simulation system contained a total of ∼156,000 atoms. The cutoff distance of van der Waals force (vdW) and short-range electrostatic interaction was set to 10 Å. Long-distance electrostatic interactions were handled using the particle net Ewald method [60]. The neighbor list of interactions was updated every five steps with a time step of 2fs. Then following procedures were then performed for running each simulation: (i) 20,000 steps energy minimization was conducted using the steepest descent algorithm; (ii) 200 ps of NVT equilibration was conducted, followed by (iii) 500 ps of NPT equilibration by position restraining the heavy atoms with a force constant of 1000 kJ mol^-1^ nm^2^. The temperature was maintained at 310 K using a velocity rescaling thermostat with a coupling constant of 0.1 ps^−1^ [61]. (iiii) Finally, a 1-μs MD simulation under NTP ensemble were conducted at 310 K and 1 atm using the velocity rescaling thermostat and the Parrinello-Rahman Barostat, respectively. [62, 63].

### Calculation of protein-DNA HBs and SBs

The HB interactions and SB interactions formed (>10%) in the last 800 ns of the simulation between the RNAP and the bound region of the promotor DNA. To determine the HBs, the distance between the donor atom and the acceptor atom is less than 3.5 Å, and the angle of the donor atom-hydrogen atom-acceptor atom is greater than 140 degrees. And the salt bridge is defined as the distance between the most positively charged N atom of the Arg or Lys residue in the protein to the most negatively charged two oxygen atoms on phosphate group of the nucleotide is less than 5 Å.

### Calculation of protein-DNA interaction energetics

The protein-DNA interactions were calculated between two residue groups: One group is the promotor DNA (ds-DNA -17 to -1), the other group is the core part of protein within 25 Å of the promoter ds-DNA (−17 to -1). The electrostatic (*ele*) and vdW interactions were re-calculated from simulated trajectories with the water-bearing model, using the g_energy module in Gromacs. The energetics between RNAP and T7/T3 promoters and their differences for the key residues (see **Fig 3** and **SI Fig S2**) are recorded in **Table S1a** and **Table S1b**. The convergences of the calculated energetics (electrostatics) with different sizes of the protein included and different simulation time used are shown in **SI FigS1**.

