## Supplementary Information for "Switching Promotor Recognition of Phage RNA Polymerase in Silico Following Path along Lab Directed Evolution"

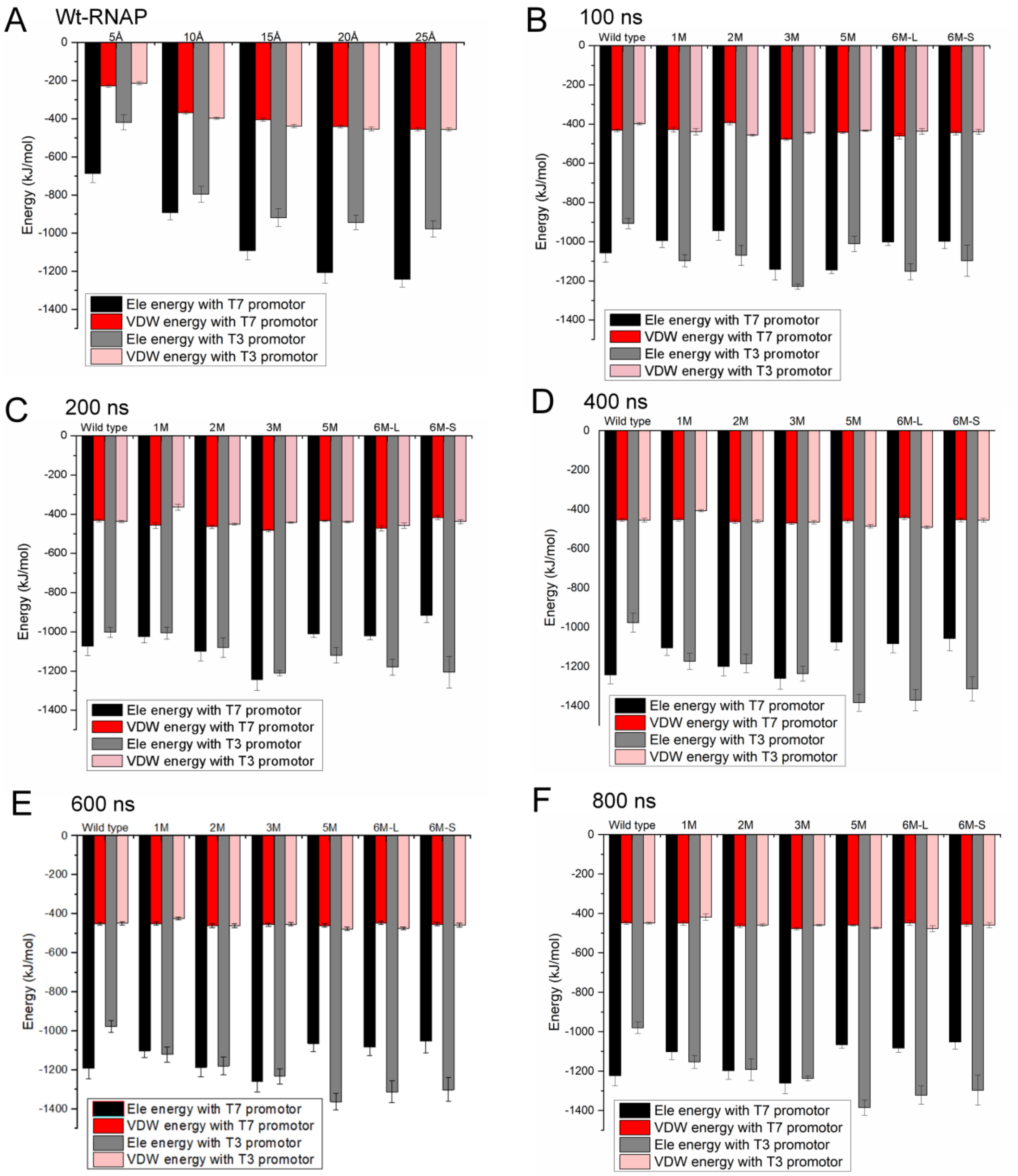


**Fig S1** The convergence of the interaction energetics. (A) Interaction energetics between the wt-RNAP and T7/T3 promoter from 5Å to 25 Å. Convergence shows after ~ 20 Å. (B-F) show the interaction energetics between the wt-RNAP and T7/T3 promoter from 100-ns to 800-ns for 14 simulation systems, i.e. wt-RNAP and six mt-RNAPs (1M to 6M), in complex with T7/T3 promotor. Convergence shows after ~ 600 ns.

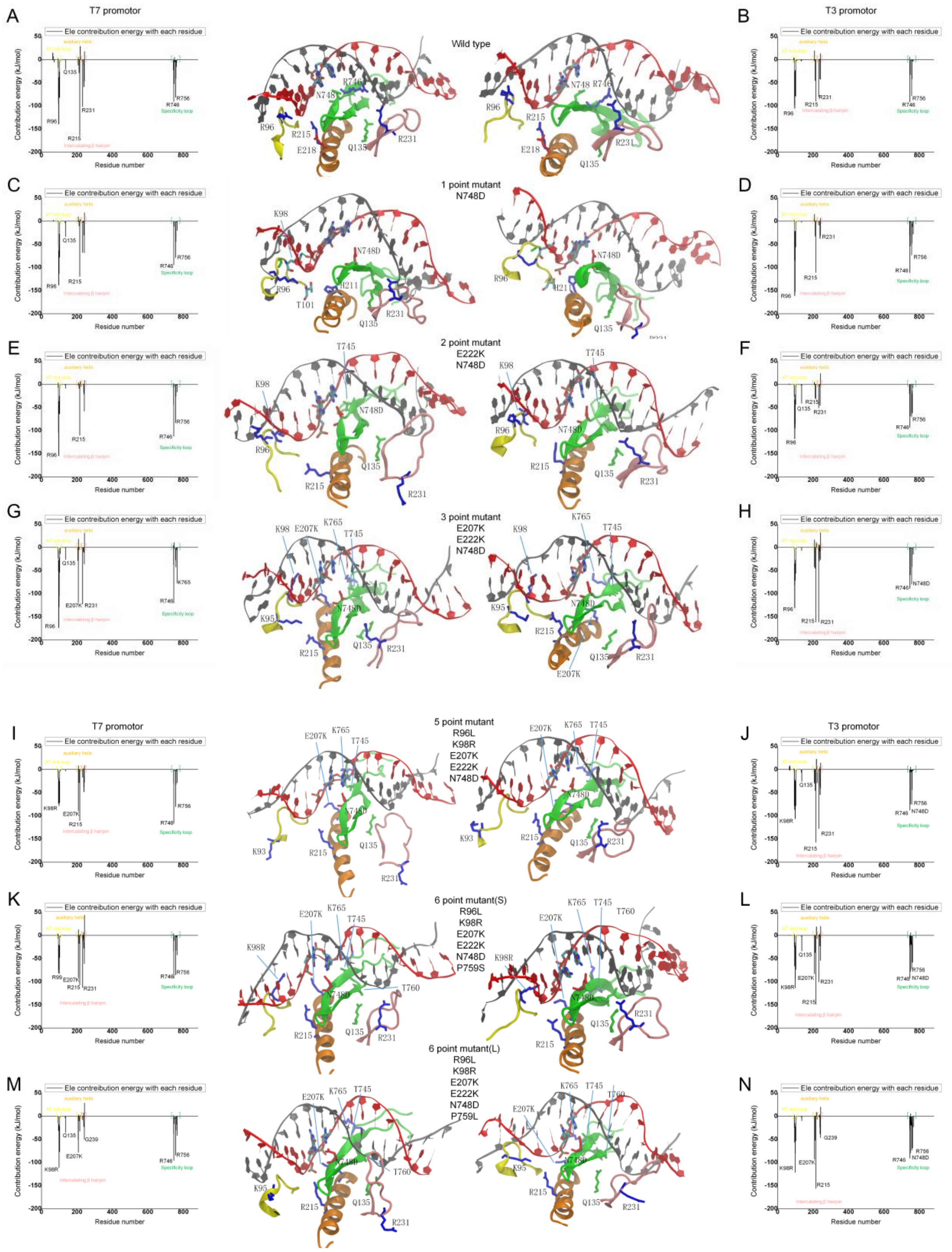


**Fig S2** The contributions from individual amino acids to electrical interaction energies in all the 14 simulation systems. The energetic contributions are denoted as $E_{T7}^{ele}$ $and$ $E_{T7}^{ele}$ in the main text for the RNAP on the T7 and T3 promoter, respectively. Key residues with notable contributions are labeled, for the wt-RNAP and six mt-RNAPs (with molecular views). See numerical values of the energetic contributions in **Table S1** below.

| WT | | WT | | 1M | | 1M | | 2M | | 2M | | 3M | | 3M | |
|---|---|---|---|---|---|---|---|---|---|---|---|---|---|---|---|
| residue id | E_T7 | residue id | E_T3 | residue id | E_T7 | residue id | E_T3 | residue id | E_T7 | residue id | E_T3 | residue id | E_T7 | residue id | E_T3 |
| 63 | 14.73 | 95 | -98.14 | 95 | -52.62 | 95 | -83.6 | 95 | -104.2 | 93 | -26.9 | 93 | -22.6 | 95 | -81.3 |
| 93 | -17.16 | 96 | -105.37 | 96 | -138.21 | 96 | -161 | 96 | -154.6 | 95 | -93.6 | 95 | -50.2 | 96 | -130.9 |
| 95 | -94.98 | 97 | -26.58 | 97 | -63.45 | 97 | -22 | 97 | -36.8 | 96 | -117.6 | 96 | -173.4 | 97 | -34.9 |
| 96 | -139.4 | 98 | -22.78 | 98 | -113.5 | 98 | -152.5 | 98 | -63.7 | 97 | -33.2 | 97 | -28.5 | 98 | -64.8 |
| 97 | -38.6 | 99 | -78.35 | 99 | -63.3 | 99 | -85.2 | 99 | -52.8 | 98 | -95.1 | 98 | -128.6 | 99 | -71.3 |
| 98 | -45.7 | 100 | -24.6 | 100 | -27.8 | 100 | -21.5 | 100 | -25.3 | 99 | -52.3 | 99 | -103.7 | 100 | -27.2 |
| 99 | -82.6 | 101 | -79.01 | 101 | -78.03 | 101 | -55.5 | 101 | -64.1 | 100 | -22.6 | 100 | -22.2 | 101 | -56.3 |
| 100 | -26.28 | 208 | 19.29 | 135 | -33.2 | 208 | 15.2 | 215 | -110.4 | 101 | -43.3 | 101 | -53.7 | 207 | -42.2 |
| 101 | -81.7 | 211 | -45.25 | 215 | -119.5 | 211 | -32.3 | 238 | -16.2 | 135 | -41.1 | 104 | -13.4 | 208 | 16.7 |
| 135 | -20.7 | 215 | -90.07 | 231 | -67.5 | 215 | -110.9 | 239 | -21 | 211 | -29.9 | 135 | -26.2 | 211 | -45.4 |
| 206 | -12.4 | 231 | -80.39 | 239 | -30.1 | 239 | -37.3 | 241 | -57.9 | 215 | -31.7 | 205 | -28.8 | 215 | -161.5 |
| 211 | -28.6 | 239 | -42.11 | 241 | -68.4 | 241 | -31.6 | 745 | -43.4 | 231 | -51.1 | 207 | -119.3 | 218 | 10.5 |
| 215 | -164.6 | 241 | -42.24 | 242 | 18.3 | 745 | -30.9 | 746 | -112.5 | 239 | -15.6 | 208 | 18.3 | 231 | -161.3 |
| 218 | 28.9 | 242 | 12.96 | 746 | -94.4 | 746 | -111.3 | 756 | -83.5 | 241 | -45.9 | 231 | -122.7 | 234 | -21.9 |
| 231 | -101.8 | 746 | -91.49 | 748 | -11 | 748 | -53.5 | 758 | -26.2 | 242 | 23.4 | 238 | -10.7 | 235 | -29.3 |
| 238 | -11.63 | 748 | -32.4 | 756 | -78.9 | 756 | -73.2 | 765 | -16.7 | 744 | -10.3 | 241 | -36.1 | 241 | -26.3 |
| 239 | -58.8 | 756 | -78.19 | 758 | -44.9 | 758 | -47.4 | | | 746 | -93.5 | 242 | 31.4 | 242 | 31.9 |
| 241 | -46.2 | 758 | -41.75 | 765 | -20.2 | 760 | -11.6 | | | 748 | -68 | 745 | -37.7 | 746 | -88.6 |
| 242 | 17.26 | | | | | 765 | -28.1 | | | 756 | -70.5 | 746 | -120.2 | 748 | -59.4 |
| 744 | -22.3 | | | | | | | | | 757 | -15.1 | 748 | 10.5 | 756 | -79.9 |
| 746 | -90.4 | | | | | | | | | 758 | -46.7 | 756 | -27.2 | 758 | -41.5 |
| 748 | -52 | | | | | | | | | 759 | -20.8 | 758 | -42.4 | 765 | -15.1 |
| 756 | -81.5 | | | | | | | | | 760 | -31.8 | 765 | -77.5 | | |
| 758 | -40.9 | | | | | | | | | | | | | | |
| 760 | -15 | | | | | | | | | | | | | | |

| 5M | | 5M | | 6M-1 | | 6M-1 | | 6M-2 | | 6M-2 | |
|---|---|---|---|---|---|---|---|---|---|---|---|
| residue id | E_T7 | residue id | E_T3 | residue id | E_T7 | residue id | E_T3 | residue id | E_T7 | residue id | E_T3 |
| 93 | -18.29 | 93 | -78.08 | 95 | -80.9 | 93 | -24 | 93 | -15.2 | 93 | -24 |
| 95 | -73.5 | 95 | -86.64 | 96 | -20 | 95 | -66.1 | 95 | -31.2 | 95 | -56.1 |
| 96 | -19.05 | 96 | -21.8 | 97 | -29.8 | 96 | -15.5 | 96 | -17.3 | 96 | -15.5 |
| 97 | -40.13 | 97 | -31.84 | 98 | -72.9 | 97 | -26.1 | 97 | -18.4 | 97 | -36.1 |
| 98 | -78.53 | 98 | -106.7 | 99 | -83.5 | 98 | -113.4 | 98 | -110.8 | 98 | -121.4 |
| 99 | -73.92 | 99 | -104.5 | 100 | -21.8 | 99 | -95.1 | 99 | -61.5 | 99 | -80.1 |
| 100 | -25.23 | 100 | -25.21 | 101 | -47.6 | 100 | -22.8 | 100 | -22.6 | 100 | -22.8 |
| 101 | -71.08 | 101 | -78.13 | 207 | -90.6 | 101 | -73.2 | 101 | -76.8 | 101 | -53.2 |
| 207 | -89.23 | 135 | -25.88 | 208 | 12.3 | 135 | -33.7 | 135 | -35.6 | 207 | -93.1 |
| 211 | -28.44 | 206 | -30.03 | 211 | -17.5 | 207 | -83.1 | 206 | -50.2 | 211 | -52.1 |
| 215 | -110.75 | 207 | -27.47 | 215 | -107.1 | 211 | -37.1 | 207 | -78.2 | 215 | -156 |
| 231 | -16.77 | 211 | -45.78 | 231 | -112.8 | 215 | -149 | 215 | -32.4 | 237 | -20.9 |
| 237 | -15.35 | 215 | -157.28 | 235 | -10.1 | 218 | 18.7 | 237 | -14.2 | 238 | -22.8 |
| 238 | -38.31 | 218 | 22.46 | 237 | -15.2 | 231 | -100.7 | 238 | -21.9 | 239 | -38.2 |
| 239 | -48.11 | 222 | -29.06 | 238 | -25.9 | 237 | -10.9 | 239 | -22.9 | 241 | -46.2 |
| 241 | -27.94 | 231 | -127.67 | 239 | -28 | 238 | -22.8 | 241 | -50.4 | 242 | 19.8 |
| 745 | -30.07 | 237 | -16.34 | 241 | -60.9 | 239 | -17.1 | 242 | 28.4 | 744 | -21.3 |
| 746 | -116.18 | 238 | -35.22 | 242 | 44.1 | 241 | -54.2 | 745 | -38.8 | 745 | -30.2 |
| 756 | -80.2 | 239 | -54.41 | 745 | -34.2 | 242 | 19.8 | 746 | -95.8 | 746 | -90.8 |
| 758 | -32.36 | 241 | -40.51 | 746 | -92.5 | 744 | -22.3 | 748 | -16 | 748 | -79.3 |
| 765 | -28.65 | 242 | 14.39 | 748 | -10.9 | 746 | -92.8 | 756 | -84.4 | 751 | -17.5 |
| | | 746 | -87.61 | 756 | -85.2 | 748 | -77.7 | 758 | -30.4 | 752 | -10.2 |
| | | 748 | -74.64 | 758 | -32.6 | 751 | -17.5 | 759 | -16.1 | 753 | -20.2 |
| | | 756 | -75.26 | 765 | -42.2 | 752 | -10.2 | 760 | -16.8 | 756 | -73.9 |
| | | 758 | -43.68 | | | 753 | -20.2 | 765 | -41.8 | 758 | -41.6 |
| | | | | | | 756 | -71.9 | | | 759 | -14.7 |
| | | | | | | 758 | -37.6 | | | 760 | -68.9 |
| | | | | | | 759 | -17.7 | | | 765 | -23.1 |
| | | | | | | 760 | -57.9 | | | | |

**Table S1a**: The significant residue contributions to the electrostatic interaction energetics between RNAPs and T7/T3 promoter ($E_{T7}^{ele}$ & $E_{T3}^{ele}$; absolute values larger than 10 kJ/mol).

| WT | | 1M | | 2M | | 3M | | 5M | | 6M-1 | | 6M-2 | |
|---|---|---|---|---|---|---|---|---|---|---|---|---|---|
| residue id | ΔE | residue id | ΔE | residue id | ΔE | residue id | ΔE | residue id | ΔE | residue id | ΔE | residue id | ΔE |
| 63 | 14.73 | 95 | 30.98 | 93 | 19.6 | 93 | -21.8 | 93 | 59.79 | 93 | 18 | 95 | 24.9 |
| 93 | -12.41 | 96 | 22.79 | 95 | -10.6 | 95 | 31.1 | 95 | 13.14 | 95 | -14.8 | 97 | 17.7 |
| 96 | -34.03 | 97 | -41.45 | 96 | -37 | 96 | -42.5 | 98 | 28.17 | 98 | 40.5 | 98 | 10.6 |
| 97 | -12.02 | 98 | 39 | 98 | 31.4 | 98 | -63.8 | 99 | 30.58 | 99 | 11.6 | 99 | 18.6 |
| 98 | -22.92 | 99 | 21.9 | 101 | -20.8 | 99 | -32.4 | 135 | 22.45 | 101 | 25.6 | 101 | -23.6 |
| 135 | -18.96 | 101 | -22.53 | 135 | 32.2 | 104 | -12.3 | 206 | 28.47 | 135 | 33.5 | 135 | -25.9 |
| 208 | -18.17 | 135 | -30.7 | 211 | 20.6 | 135 | -19.1 | 207 | -61.76 | 208 | 11.3 | 206 | -50.2 |
| 211 | 16.65 | 208 | -11.92 | 215 | -78.7 | 205 | -28.8 | 211 | 17.34 | 211 | 19.6 | 207 | 14.9 |
| 215 | -74.53 | 211 | 22.33 | 231 | 50.6 | 207 | -77.1 | 215 | 46.53 | 215 | 41.9 | 211 | 42.13 |
| 218 | 28.844 | 231 | -59.3 | 238 | -13.9 | 211 | 39.9 | 218 | -22.46 | 218 | -16.3 | 215 | 123.6 |
| 231 | -21.41 | 241 | -36.8 | 241 | -12 | 215 | 161.5 | 222 | 28.67 | 231 | -12.1 | 239 | 15.3 |
| 239 | -16.69 | 242 | 11.8 | 242 | -15.1 | 218 | -10.5 | 231 | 110.9 | 239 | -10.9 | 744 | 23.4 |
| 744 | -28.19 | 745 | 29.1 | 744 | 12.2 | 231 | 38.6 | 241 | 12.57 | 242 | 24.3 | 748 | 63.3 |
| 748 | -19.6 | 746 | 16.9 | 745 | -43.4 | 233 | -15.5 | 745 | -29.7 | 744 | 26.9 | 750 | -11.7 |
| 760 | -15.77 | 748 | 42.5 | 746 | -19 | 234 | 21.6 | 746 | -28.57 | 745 | -34.1 | 751 | 17.7 |
| | | 760 | 10.1 | 748 | 63.1 | 235 | 29 | 748 | 71.79 | 748 | 66.8 | 753 | 19.5 |
| | | | | 756 | -13 | 745 | -36.8 | 758 | 11.32 | 751 | 17.6 | 756 | -10.5 |
| | | | | 757 | 14.2 | 746 | -31.6 | 765 | -27.92 | 752 | 10.1 | 758 | 11.2 |
| | | | | 758 | 20.5 | 748 | 69.9 | | | 753 | 19.5 | 760 | 52.1 |
| | | | | 759 | 19.6 | 756 | 52.7 | | | 756 | -13.3 | 765 | -18.7 |
| | | | | 760 | 29.9 | 765 | -62.4 | | | 759 | 16.8 | | |
| | | | | 765 | -16.2 | | | | | 760 | 54.3 | | |
| | | | | | | | | | | 765 | -42.1 | | |

**Table S1b**: The significant residue contributions to the RNAP electrostatic energetics differences between T7 and T3 promoters ($\Delta E^{ele} = E_{T7}^{ele} - E_{T3}^{ele}$; absolute values larger than 10 kJ/mol).

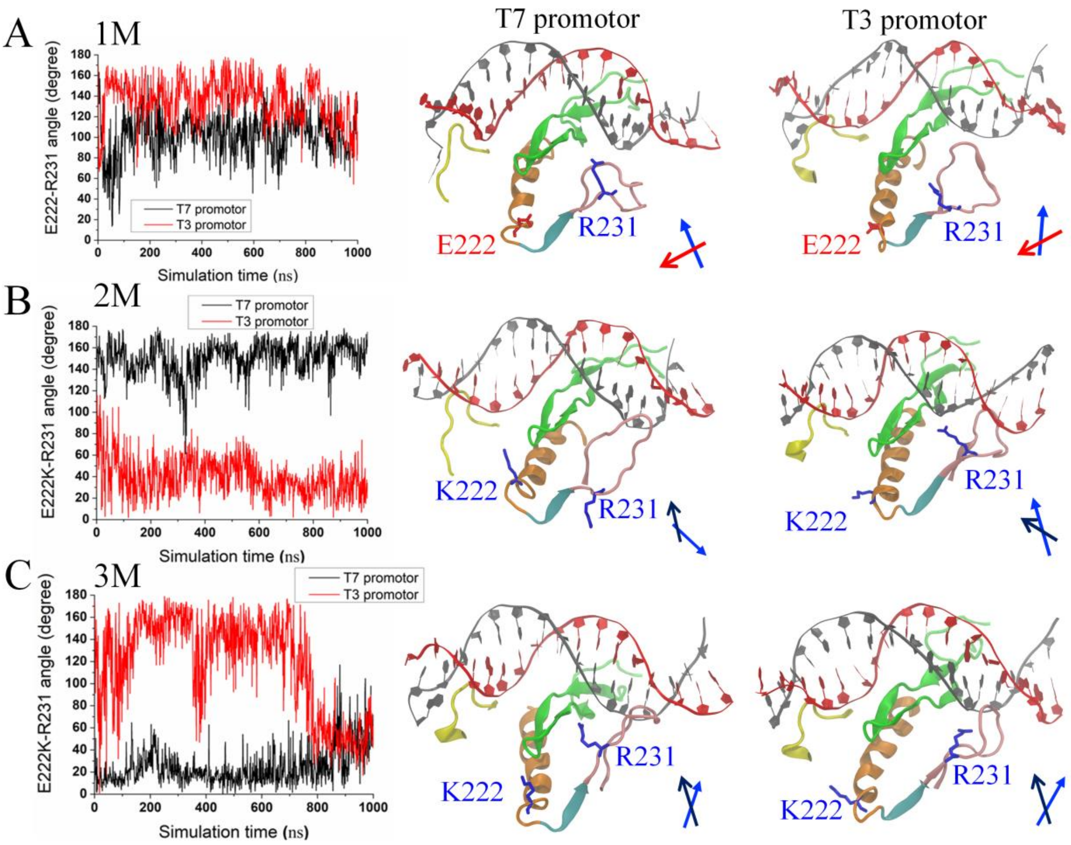


**Fig S3** Structural views on re-orientations between E222K and R231 for three RNA variants (1M to 3M) in complexes with T7 /T3 promoter, respectively. The orientational angle (θ) between the side chain of E222K and the side chain of K231 are measured from the simulations (*left*; with black/red data for T7/T3 promoter system). In the molecular views (*right*), the AT-rich loop (ATL, in yellow), specificity loop (SPL, green), intercalating β hairpin (INB, pink) and auxiliary helix (AXH, in orange) that are in close association with the DNA promoter are shown. A cartoon on the orientation angle between E222K and R231 is shown (*bottom right*) for each simulation system/molecular view.

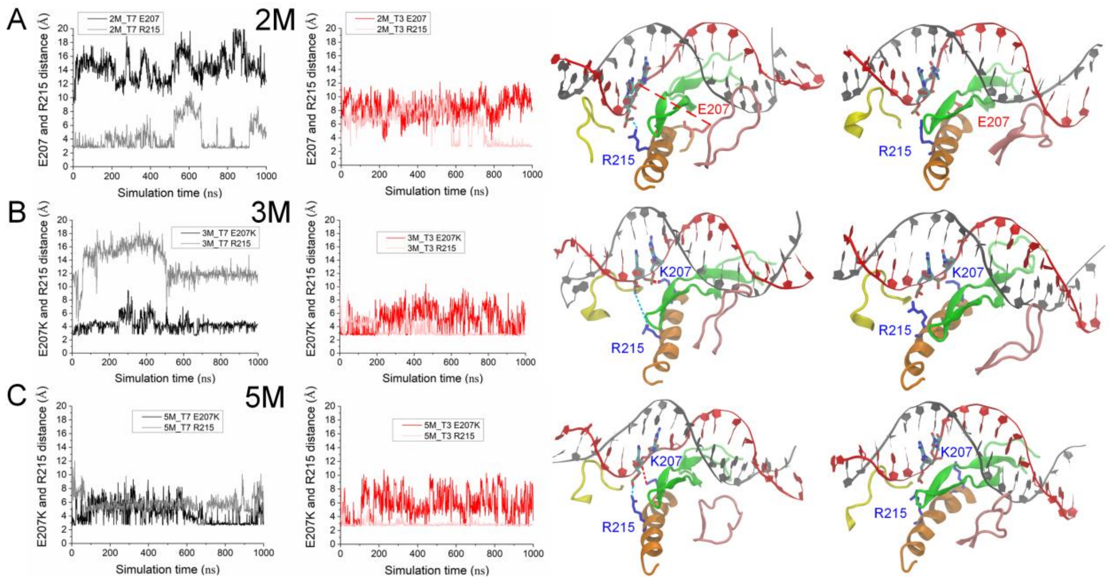


**Fig S4** Structural views of potential hydrogen bond (HB) interactions between E207K/R215 and DNA promotor for wt-RNAP and three RNA variants (2M, 3M and 5M) in complexes with T7 /T3 promoter, respectively. Two distances for potential HB between E207K and G-11 and HB between R215 and C-12 are measured from the simulations (*left*, with black/red data for T7/T3 promoter system). In the molecular views (*right*), the AT-rich loop (ATL, in yellow), specificity loop (SPL, green), intercalating β hairpin (INB, pink) and auxiliary helix (AXH, in orange) that are in close association with the DNA promoter are shown, with E207 or K207, R215 and G-11, C-12 on the non-template strand labeled.

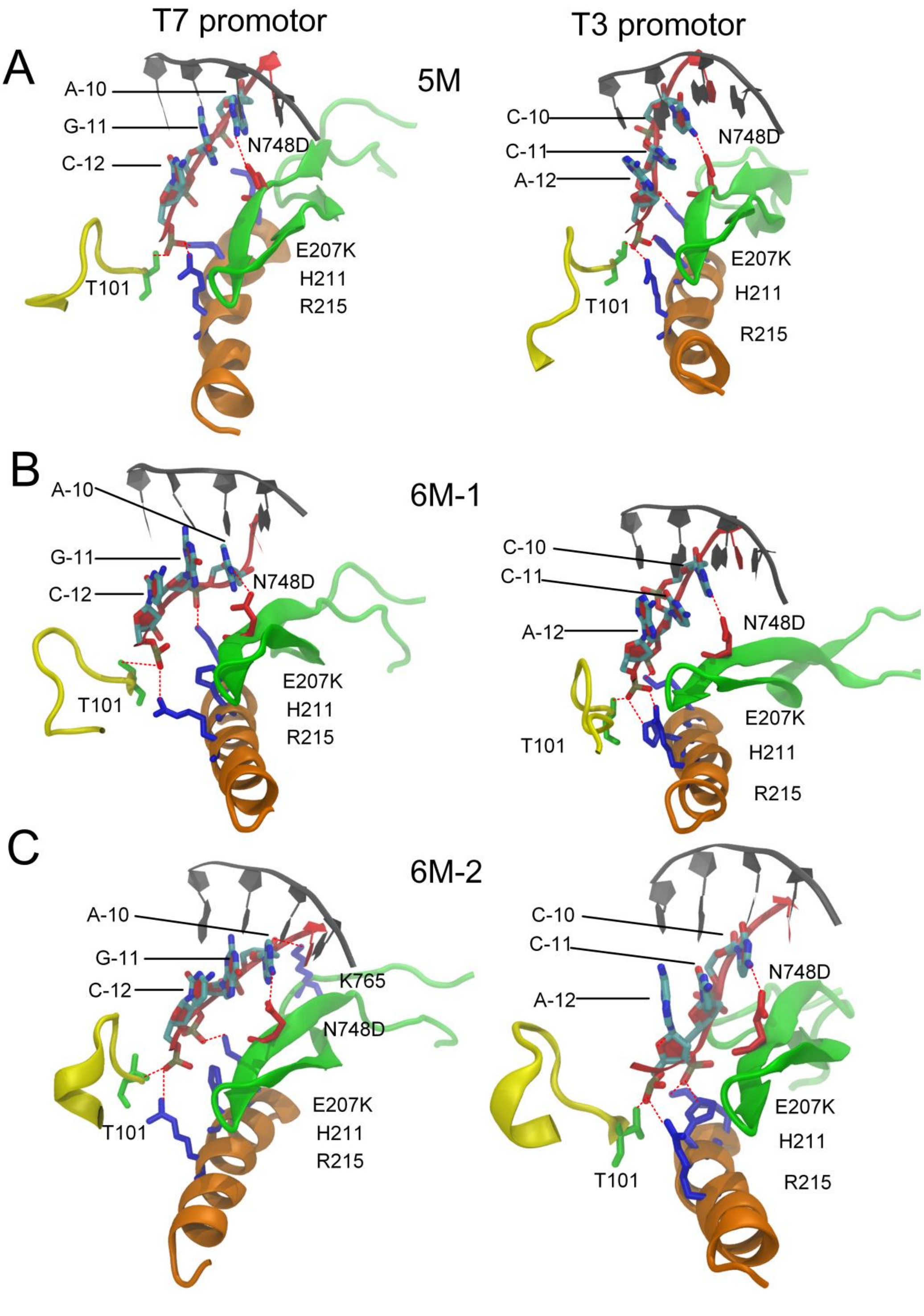


**Fig S5**. The hydrogen bonds (HBs) between the RNAP and the central bound region (-12 to -10) of promotor DNA. The simulations systems are shown for the directed mutants 5M (A), 6M-1 (B), and 6M-2 (C), on the T7 (*left*) and T3 (*right*) promoter.

| | | | -15 | | | | | -10 | | | | | -5 |
|---|---|---|---|---|---|---|---|---|---|---|---|---|---|
| T7 template | A | T | T | A | T | G | C | T | G | A | G | T | G |
| T7 non-template | T | A | A | T | A | C | G | A | C | T | C | A | C |
| T3 non-template | A | A | T | T | A | A | C | C | C | T | C | A | C |
| | DNA bound to Protein | | | | | | | | | | | | |
| **W_T7** | R99-T<br>R96-NT | R99-T<br>R96-NT | R96-NT<br>K98-T | K95-NT<br>R96-NT<br>K93-NT | K95-NT<br>R215-NT | R215-NT | | | | R231-T | R746-T<br>R231-T | | |
| **W_T3** | R99-T<br>R93-T<br>R96-NT | R99-T<br>R96-NT<br>K98-T | R96-NT<br>K98-T | K95-NT<br>R96-NT | K95-NT<br>R215-NT | R215-NT | | | | R231-T | R746-T<br>R231-T | R746-T | |
| **M1_T7** | R99-T<br>R96-NT | R99-T<br>R96-NT | R96-NT | K95-NT<br>R96-NT | K95-NT<br>R215-NT | R215-NT<br>K98-NT | K98-NT | K765-NT | | R231-T | R746-T<br>R231-T | | |
| **M1_T3** | R99-T<br>R96-NT | R99-T<br>R96-NT<br>K98-NT | R96-NT<br>R99-T | K95-NT<br>K98-T | K95-NT<br>R215-NT | R215-NT<br>K95-NT | K98-NT<br>R215-NT | K98-NT | | R231-T | R746-T | | |
| **M2_T7** | R99-T | R99-T<br>K98-T<br>R96-NT | R96-NT<br>K98-T | K95-NT<br>R96-NT<br>K93-NT | K95-NT<br>R99-NT<br>R215-NT | R215-NT<br>K98-NT | K98-NT | | | | R746-T | | |
| **M2_T3** | R99-T<br>R93-T | R99-T<br>K98-T | R96-NT<br>K98-T | K95-NT<br>R96-T | K95-NT<br>R215-NT | R215-NT<br>K98-NT | | | | R231-T | R746-T<br>R231-T | R746-T | |
| **M3_T7** | R99-T<br>R96-NT | R99-T<br>R96-NT | R96-NT | K98-T | K95-NT | K95-NT | K207-NT | K207-NT | | R231-T<br>R756-T | R231-T | R746-T | |
| **M3_T3** | R99-T<br>R93-T<br>R96-NT | R99-T<br>R96-NT<br>K98-T | R96-NT<br>K98-T | R96-NT<br>K95-NT | K95-NT<br>R215-NT | R215-NT<br>K98-NT | K207-NT | K207-NT | | | R231-T | R231-T | |
| **M5_T7** | R99-T<br>K93-T | R99-T | R98-T | R95-NT | K95-NT<br>R215-NT | R98-NT<br>R215-NT | R98-NT<br>K207-NT | K207-NT<br>K765-NT | | | R746-T<br>R231-T | R746-T | |
| **M5_T3** | R99-T<br>K93-T | R99-T<br>R98-T | R98-T | K95-NT<br>K222-NT | K95-NT<br>R215-NT<br>K222-NT | R98-NT<br>R215-NT | R98-NT<br>K207-NT | | | | R231-T | R231-T | |
| **M6(L)_T7** | R99-T<br>K93-T | R99-T | R98-T | R95-NT | R95-NT<br>R99-NT<br>R215-NT | R98-NT<br>R215-NT | R98-NT<br>K207-NT | K207-NT | K765-NT | R231-T | R231-T | | |
| **M6(L)_T3** | R99-T<br>K95-T<br>K93-T | R99-T<br>K93-NT<br>R98-T | R98-T | | K95-NT<br>R215-NT<br>K222-NT | R98-NT<br>R215-NT | R98-NT<br>K207-NT | | R231-T | R231-T | R231-T | R746-T | |
| **M6(S)_T7** | R99-T<br>K93-T | R99-T<br>R98-T | R98-T | R95-NT | R95-NT<br>R215-NT | R98-NT<br>R215-NT | K207-NT | K207-NT<br>K765-NT | | | | | |
| **M6(S)_T3** | R99-T<br>K93-T | R99-T<br>R98-T | R98-T | R95-NT | K95-NT<br>K222-NT | R98-NT<br>R215-NT | R98-NT<br>K207-NT | | K765-NT | R231-T | R231-T | | |

**Fig S6**. Analysis of salt-bridge interaction between RNAP and the promotor DNA in all simulation systems. The color convention and labelling are the same as main **Fig 5A**.

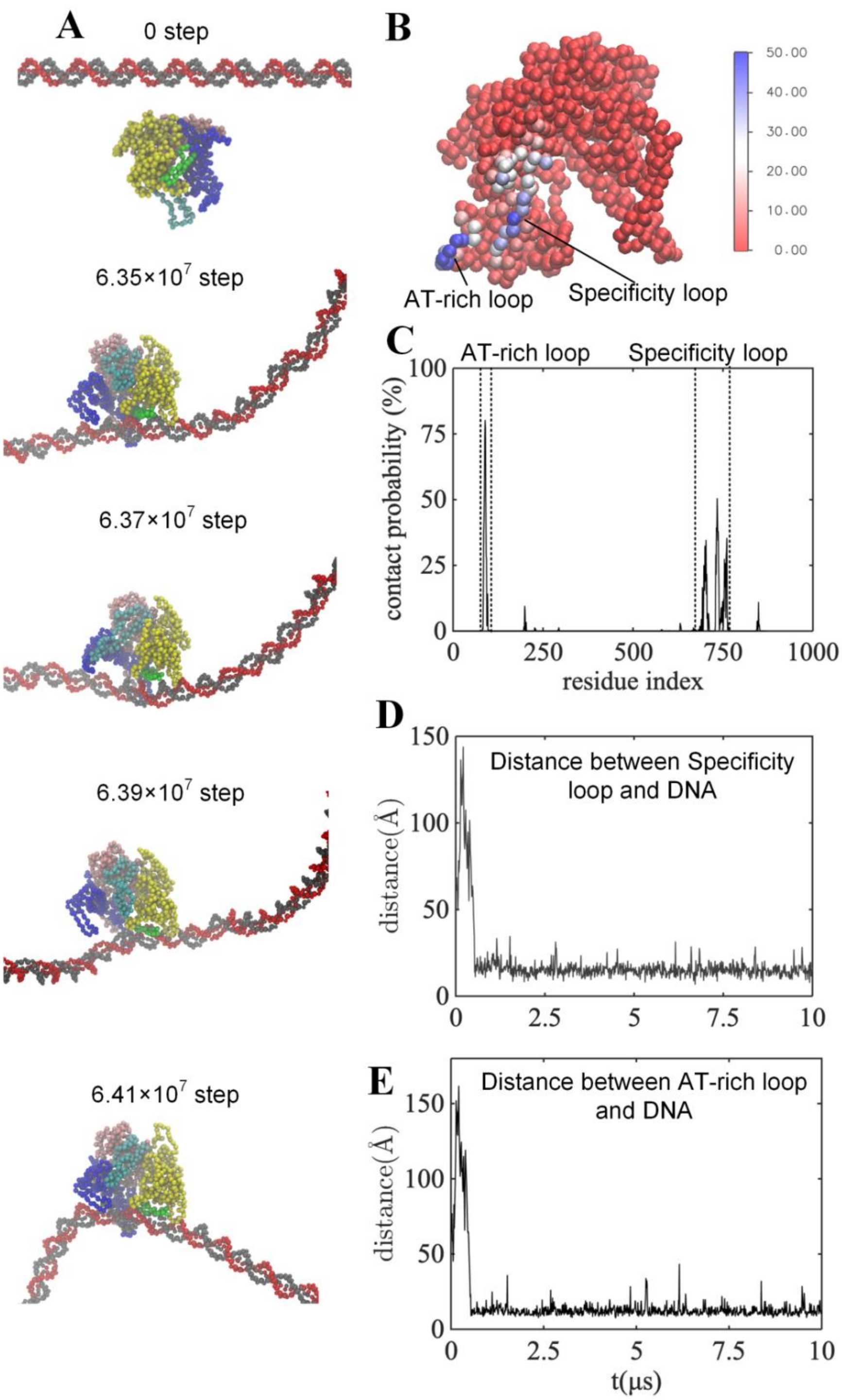


**Fig S7**. Coarse-grained (CG) modeling and MD simulation of wt-T7 RNAP diffusional search along DNA. (A) Different conformations/snapshots of the T7 RNAP on the DNA captured in the CG simulation. The N-terminal, thumb, palm, and fingers subdomains of the RNAP structure are colored by yellow, cyan, pink, and blue, respectively. The specificity loop (SPL, residue 739-770 are shown in green). The ds-DNA are shown in gray and red strands. (B and C) Contact probability of T7 RNAP and DNA, mapped in the RNAP structure (B) and shown in the diagram for all residues (C). (D and E) The distance between mass center of specificity loop (D) or AT-rich loop (E) and the nearest DNA nucleotide.

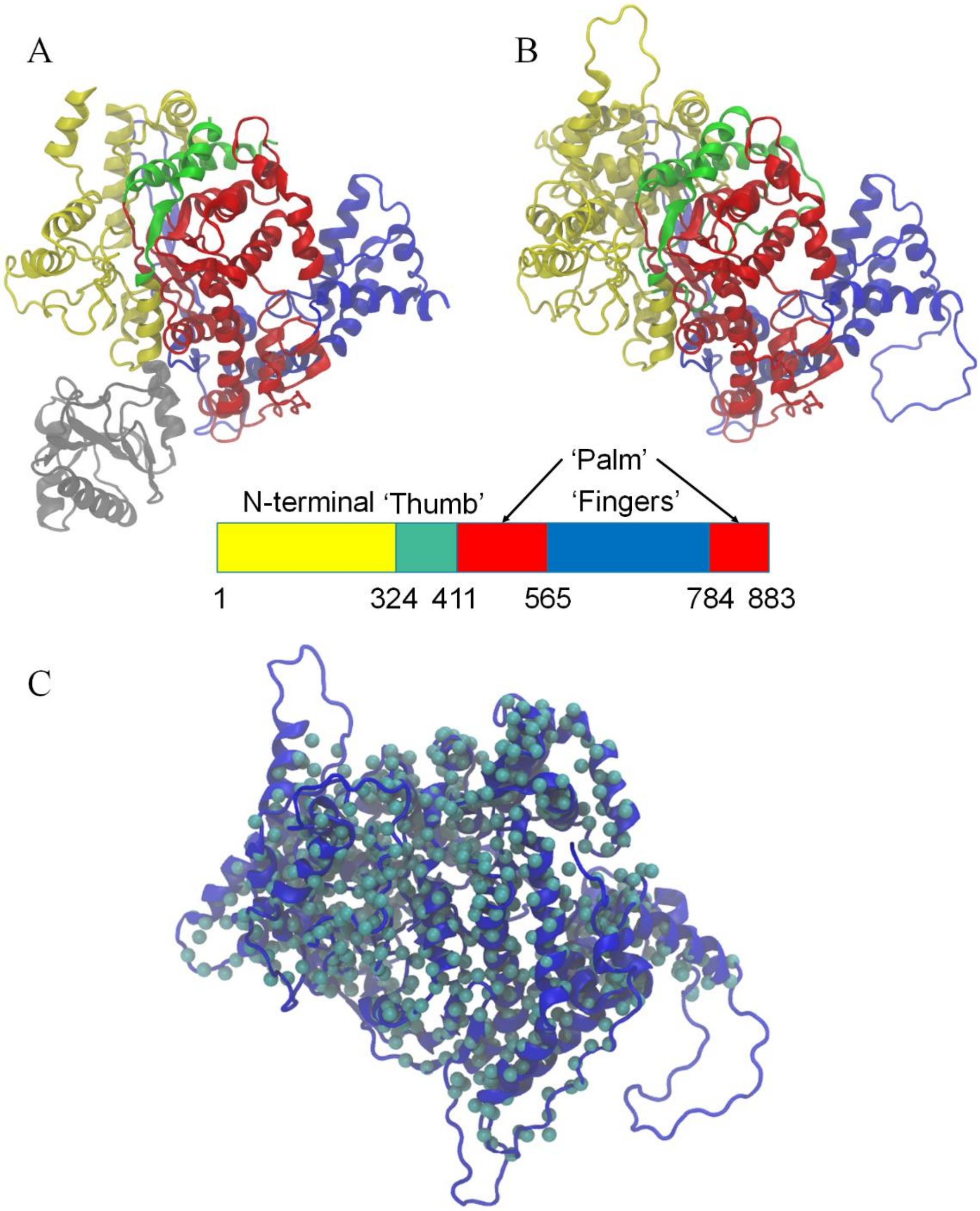


**Fig S8**. Molecular views of individual T7 RNAP structures obtained from the crystal structure studies. (A) The crystal structure of T7 RNAP combined with T7 lysozyme (PD: 1ARO). (B) The T7 RNAP structure with T7 lysozyme removed and homology modeling used to fill in the missing structure. The N-terminal, thumb, palm, and fingers domain of the RNAP structure are represented by yellow, green, red, and blue, respectively. T7 lysozyme is shown in gray. (C) The comparison between the T7 RNAP structure we constructed (from B) and the crystal structure containing only Cα atoms (PDB id: 4RNP) solved by Sousa et al [123]. The overall RMSD value is ~2.9 Å, and the most significant difference is the N-terminal part (with the separately calculated RMSD reaching ~ 8.8 Å).

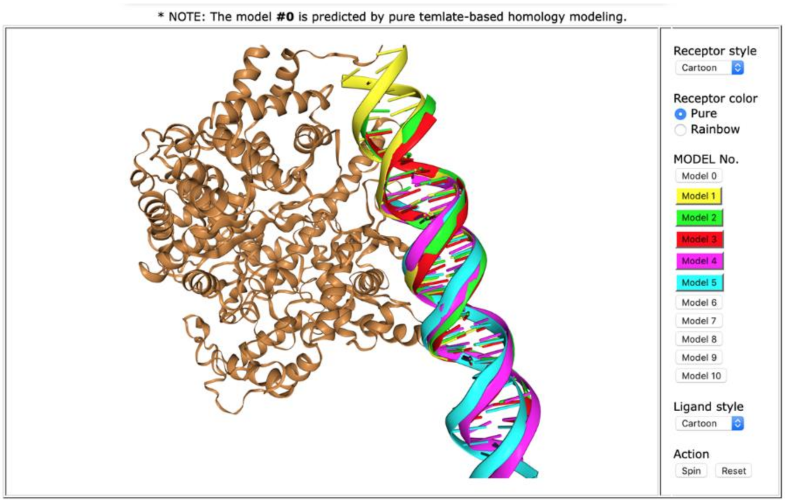


**Summary of the top 10 models**

| Rank | 1 | 2 | 3 | 4 | 5 | 6 | 7 | 8 | 9 | 10 |
|---|---|---|---|---|---|---|---|---|---|---|
| Docking Score | -548.90 | -297.61 | -278.26 | -276.91 | -264.40 | -246.21 | -246.02 | -239.03 | -237.92 | -237.79 |
| Ligand rmsd (Å) | 0.37 | 43.22 | 44.69 | 27.11 | 29.89 | 10.85 | 42.33 | 41.67 | 51.73 | 17.83 |

(a) Row 1: The ranks of the models.
(b) Row 2: The docking energy scores.
(c) Row 3: The ligand RMSDs from the input structures or modeled structures by homology modeling.

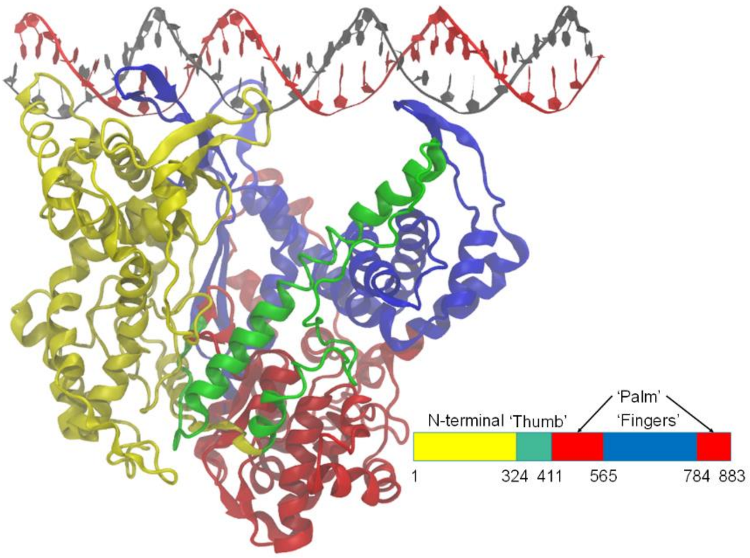


**Fig S9**. Using Hdock to construct closed complex from double-stranded DNA and T7 RNAP. The top ten structure diagram and scores selected by Hdock software in the left figure. The complex model we finally selected in the right figure, the N-terminal, Thumb, palm and fingers domain in T7 RNAP are represented by yellow, green, red and blue, respectively.

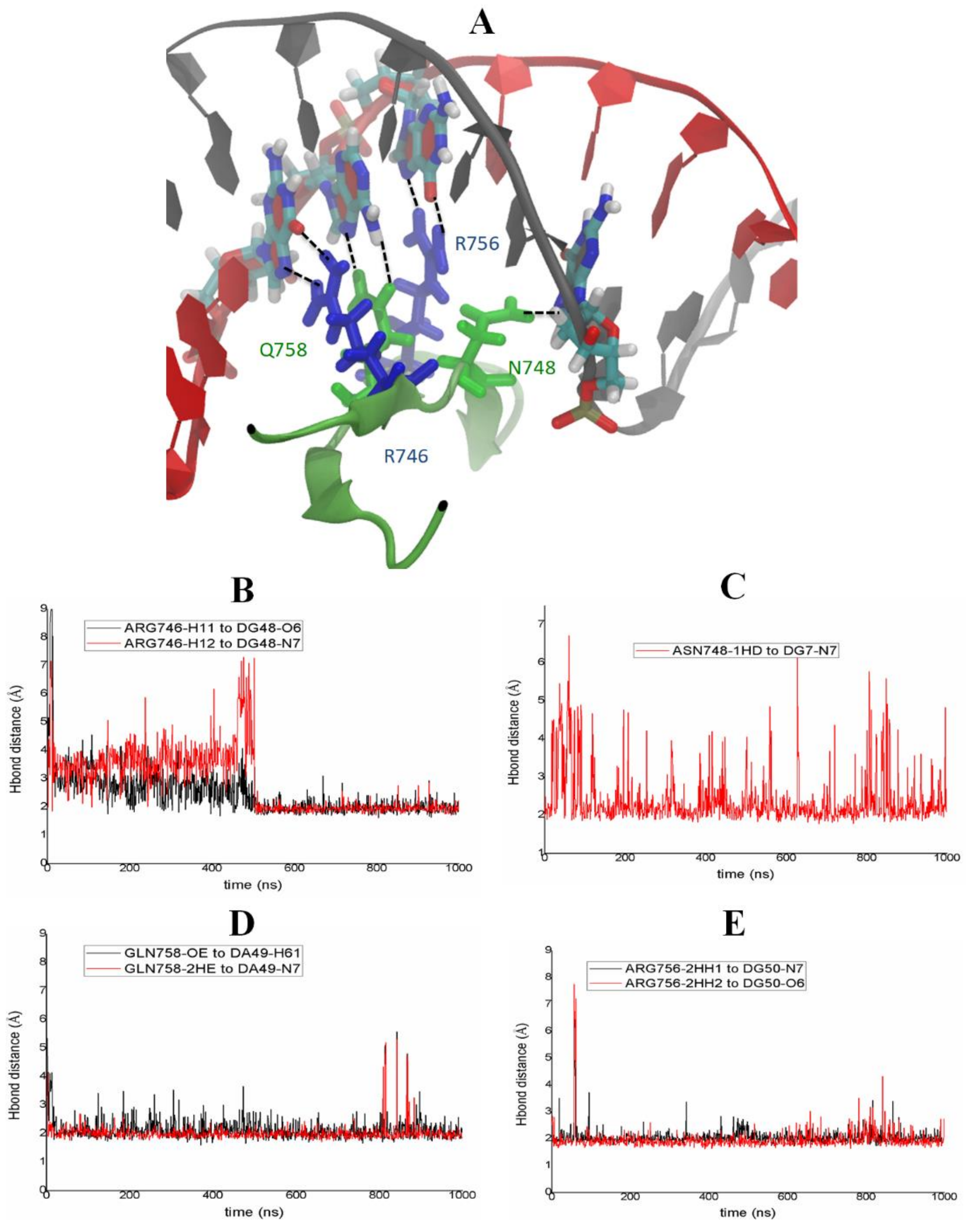


**Fig S10**. Examining hydrogen bonding interactions between the specificity loop and promotor DNA in the constructed model of the T7 RNAP initiation complex (closed). (A) structure view of the 7 hydrogen bonds formed between the four amino acids (R746, N748, R756, and Q758) and promotor DNA in the 1 us MD simulation of the selected initiation complex model. The distances corresponding to the expected hydrogen bonds are shown in (B-E).
